# Plasma Membrane Remodelling in GM2 Gangliosidoses Drives Synaptic Dysfunction

**DOI:** 10.1101/2024.08.20.608810

**Authors:** Alex S. Nicholson, David A. Priestman, Robin Antrobus, James C. Williamson, Reuben Bush, Henry G. Barrow, Emily Smith, Kostantin Dobrenis, Nicholas A. Bright, Frances M. Platt, Janet E. Deane

## Abstract

Glycosphingolipids (GSL) are important bioactive components of cellular membranes. Complex GSLs, containing sialic acid residues are known as gangliosides and are highly abundant in the brain. Diseases of ganglioside metabolism often result in severe, early-onset neurodegeneration. The ganglioside GM2 is the substrate of the hydrolytic lysosomal β- hexosaminidase A (HexA) enzyme and when subunits of this enzyme are non-functional, GM2 lipid accumulates in cells leading to the GM2 gangliosidoses, Tay-Sachs and Sandhoff diseases. We have developed high-quality i3Neuron-based models of Tay-Sachs and Sandhoff diseases, that demonstrate storage of GM2, formation of membrane whorls and accumulation of endolysosomal proteins consistent with disease phenotypes. Importantly, in addition to lysosomal dysfunction, the composition of the plasma membrane (PM) is significantly impacted in these diseases with changes in the abundance of both lipids and proteins. The changes to the PM proteome are driven in part by exocytosis of lysosomal material resulting in the aberrant accumulation of lysosomal proteins and lipids on the cell surface. The altered abundance of GM2 at the PM was striking, bringing the abundance of this precursor lipid up to that of the common neuronal gangliosides. Furthermore, the PM profiling identifies significant changes in synaptic protein abundances with direct functional impact on neuronal activity including rapid electrical firing consistent with neuronal hyperactivity. This work provides mechanistic insights into neuronal dysfunction in the GM2 gangliosidoses and highlights that these are also severe PM disorders. This work has broad implications for other lysosomal storage disorders and late-onset neurodegenerative diseases involving sphingolipid dysregulation.

## INTRODUCTION

Glycosphingolipids (GSL) are enriched in the outer leaflet of the plasma membrane (PM) where they play crucial roles in cell signalling, the immune response and neuronal function [1,2]. GSLs consist of a membrane-embedded ceramide backbone and glycosylated headgroups ranging from simple, single sugar moieties to larger, branched complex glycans containing different sugar moieties. Complex GSLs containing one or more sialic acids, are known as gangliosides and are highly abundant in the brain [3]. GSLs interact with cholesterol and proteins to form dynamic membrane microdomains that contribute to membrane organisation, receptor clustering and vesicle trafficking [4–7]. The specific repertoire and abundance of GSLs in the PM modulates membrane properties and contributes to cell identity [8,9].

The metabolism of GSLs is spatially separated in the cell, with synthesis occurring in the endoplasmic reticulum and Golgi compartments, and degradation occurring in the late endosomal and lysosomal compartments [10,11]. GSL degradation occurs sequentially, with one sugar moiety removed at a time through the action of a series of degradative enzymes. When this process is dysregulated, it causes a range of severe diseases, most of which involve neurodegeneration [3,12,13].

The enzyme β-hexosaminidase A (HexA) is a glycosyl-hydrolase that resides in the lysosome and removes the terminal N-acetyl galactosamine from the ganglioside GM2 (**Fig. 1A**), to yield the ganglioside GM3 [14,15]. The HexA enzyme is a heterodimer of alpha and beta subunits produced by two closely related genes, *HEXA* and *HEXB*, which can only process GM2 ganglioside *in vivo* when presented by the GM2 activator protein (GM2ap) (**Fig. 1B**) [16,17]. The inability to degrade GM2 causes the severe inherited disorders Tay-Sachs and Sandhoff diseases when *HEXA* or *HEXB* genes are mutated respectively, as the residual homodimers known as HexS and HexB are unable to hydrolyse GM2 [18]. Due to the high abundance of gangliosides in neuronal cells, the accumulation of undegraded GM2 within these cells leads to severe neurological dysfunction, neurodegeneration and premature death. Disease severity and age-of-onset is directly correlated with residual enzyme activity, with the most severe forms having effectively no enzyme activity resulting in death by 2 years of age [18,19].

**Figure 1.**
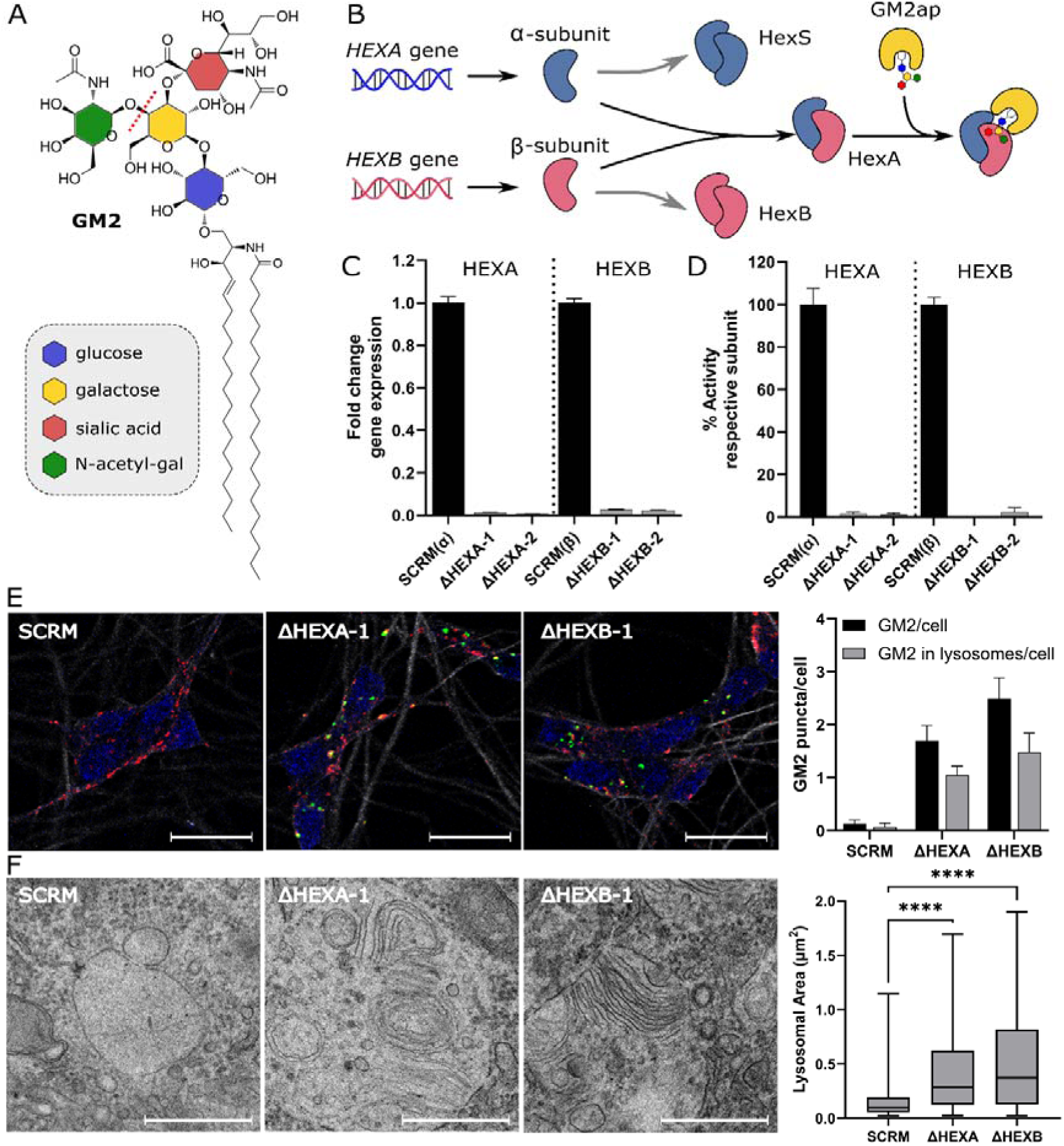
Neuronal i3N models of Tay-Sachs and Sandhoff diseases. **A.** Schematic diagram of the GM2 ganglioside detailing the composition of the glycan headgroup and illustrating which bond is cleaved by the HEXA enzyme (red dotted line). **B.** Schematic diagram of how the α- and β-subunits can form homo- and heterodimers with the HexA heterodimeric isoenzyme being the only one that can cleave GM2. GM2 is presented to HexA by the GM2 activator protein (GM2ap). **C.** Quantitative PCR (qPCR) analysis of *HEXA* and *HEXB* gene expression in neurons following CRISPRi-induced knockdown. Fold change relative to SCRM controls are shown for each cell line, n=3 biological replicates were carried out in technical triplicate and the mean is displayed ± SEM. **D.** Activity assays from cell lysates of SCRM and HEXA and HEXB CRISPRi cell lines. Relative activity of each subunit was determined by differential activity against MUG and MUGS substrates, the mean is displayed ± SEM, n=3 biological replicates were carried out and significance was calculated using a one-way ANOVA test, \*\*\*\**p* ≤ 0.0001. **E.** Fluorescence microscopy images of neurons stained for β3-tubulin (white), LAMP1 for lysosomes (red), GM2 (green) and DAPI (blue). GM2 positive puncta were quantified and analysed for co-localisation with lysosomes. Sixty images were analysed per cell line across three biological replicates. Scale bar (white line) represents 20 μm. **F.** Transmission electron microscopy images of SCRM and HEXA and HEXB CRISPRi cell lines. Zoomed images are shown to illustrate membrane whorls and zebra bodies in enlarged lysosomes. Scale bar (white line) represents 500 nm. Twenty images were analysed per cell line. Only endolysosomes where the entire compartment was visible were quantified. Significance was calculated using a one-way ANOVA test, \*\*\*\**p* ≤ 0.0001.

Tay-Sachs and Sandhoff diseases are classified as lysosomal storage diseases (LSDs) due to the presence of enlarged endolysosomes filled with undigested material [20]. In cells with a high abundance of gangliosides, such as neurons, the endolysosomes are filled with membranous substructures, referred to as whorls or zebra bodies [21]. As endolysosomes are the terminal degradative organelle for both the endocytic and autophagic pathways, defects in their homeostasis have wide reaching effects. The molecular mechanisms that link lysosomal storage to neurodegeneration are incompletely understood but lysosomal dysfunction is a common feature of many neurodegenerative conditions [22].

Several different models of Tay-Sachs and Sandhoff disease already exist, including naturally occurring cat, dog and sheep models as well as genetically modified mouse models [23]. Interestingly, mice possess some differences in GSL metabolism compared to humans, as mice have higher levels of neuraminidase that can process GM2 such that both the HEXA and NEU3 genes must be knocked out to make a Tay-Sachs model [24,25]. The differences in GSL metabolism between mice and humans supports the need to develop human-based models of these diseases for molecular insights into human pathology. iPSC-based neurons and neural progenitor cells have been developed from patient fibroblasts and exhibit lysosomal disturbances and accumulation of GM2, as well as alterations in synaptic exocytosis [26,27]. These models have been effectively used to test substrate reduction therapy and to probe disease mechanisms. However, the long differentiation time and genetic variability limit the usefulness of these models for detailed molecular analysis.

Advances in human iPSC technologies allow for new models to probe molecular mechanisms driving disease phenotypes. The i3Neuron (i3N) model is an iPSC line with a stably integrated, doxycycline-inducible NGN2 transcriptional factor for rapid differentiation into a homogenous, isogenic population of cortical glutamatergic neurons over the course of 14 days [28]. A derivative of this line also possesses a catalytically dead Cas9 (dCas9) enzyme for CRISPR interference (CRISPRi)-mediated knockdown of gene expression [29]. This i3N system allows for the rapid development of neuronal disease models that can be grown on a scale compatible with mass spectrometry based proteomic approaches.

Here we present i3N-based models for Tay-Sachs and Sandhoff diseases, which mimic disease phenotypes including enlarged endolysosomal compartments containing excessive GM2 and membrane whorls. Proteomic analysis identified significant molecular changes, including accumulation of endolysosomal proteins involved in lipid processing as well as several membrane trafficking molecules. These changes result in abnormal accumulation of lipids and proteins at the cell surface impacting neuronal functions and leading to dysregulation of electrical activity. These neuronal models of GM2 gangliosidoses represent valuable systems for undesrtanding the molecular mechanisms driving disease pathogenesis.

## RESULTS

### Accumulation of GM2 in endolysosomes of i3N models of Tay-Sachs and Sandhoff diseases

Severe, early-onset forms of Tay-Sachs and Sandhoff diseases possess less than 5% enzyme activity due to large gene deletions or missense mutations [18,19,30]. To develop relevant disease models, efficient CRISPRi-mediated knock down of *HEXA* or *HEXB* gene expression, to remove functional HEXA enzyme, was required. CRISPRi guides targeting the *HEXA* or *HEXB* genes were introduced into the dCas9 derivative of the i3N stem cells (**Supp. Table S1**). Five cell lines using different guide sequences were made from this parent dCas9 cell line, ΔHEXA-1 and ΔHEXA-2 to model Tay-Sachs disease, ΔHEXB-1 and ΔHEXB-2 to model Sandhoff and a scrambled (SCRM) non-targeting guide to be used as a control. Quantitative PCR (qPCR) analysis of neurons at the reported mature timepoint of 14 days post doxycycline-induced expression (dpi) of the NGN2 gene confirmed highly efficient knockdown of *HEXA* and *HEXB* gene expression. mRNA levels in all cell lines were between 0.5-3% of gene expression compared with SCRM lines (**Fig. 1C**). Enzyme activity assays are used to screen carriers and identify disease severity, and here they were adapted to determine activity in lysates of differentiated neurons [31,32]. This confirmed that the loss of gene expression translated into an equivalent lack of functional HEXA subunits, HEX-α and β, in the respective cell lines (**Fig. 1D**). The levels of enzyme activity shown are consistent with those seen in patients with infantile or juvenile forms of early-onset Tay-Sachs and Sandhoff diseases [18].

The most striking phenotype in cells with HEXA dysfunction is lysosomal enlargement caused by the storage of undigested GM2 lipid substrate. Immunofluorescence microscopy using an α-GM2 antibody showed accumulation of GM2 in ΔHEXA-1 and ΔHEXB-1 cell lines. The majority of this ganglioside colocalised with LAMP1 positive structures, whilst very little GM2 was detected in the SCRM line (**Fig. 1E**). Analysis of the ultrastructure of ΔHEXA-1 and ΔHEXB-1 cells using transmission electron microscopy revealed significant enlargement of endolysosomal compartments and accumulation of membranous substructures in the form of whorls and zebra bodies, absent in the SCRM line (**Fig. 1F**). This confirmation that GM2 accumulates in endolysosomes and that these compartments contain large amounts of multi-lamellar membranous structures similar to those observed in human and animal disease cells [20,21,23,33,34] supports the validity of these i3N models of Tay-Sachs and Sandhoff diseases.

Previously published work has confirmed that i3N cells express markers of functionally mature neurons by 14 dpi [28]. qPCR analysis of the i3N cell lines confirmed expression of neuronal markers (synaptophysin SYP, MAP2 and β3-tubulin) and loss of stem cell markers (NANOG and OCT-4) at this timepoint (**Fig. 2A** and **Supp. Fig. S1**). These data confirm that the disease model lines can still differentiate normally and express relevant markers of neuronal maturity.

**Figure 2.**
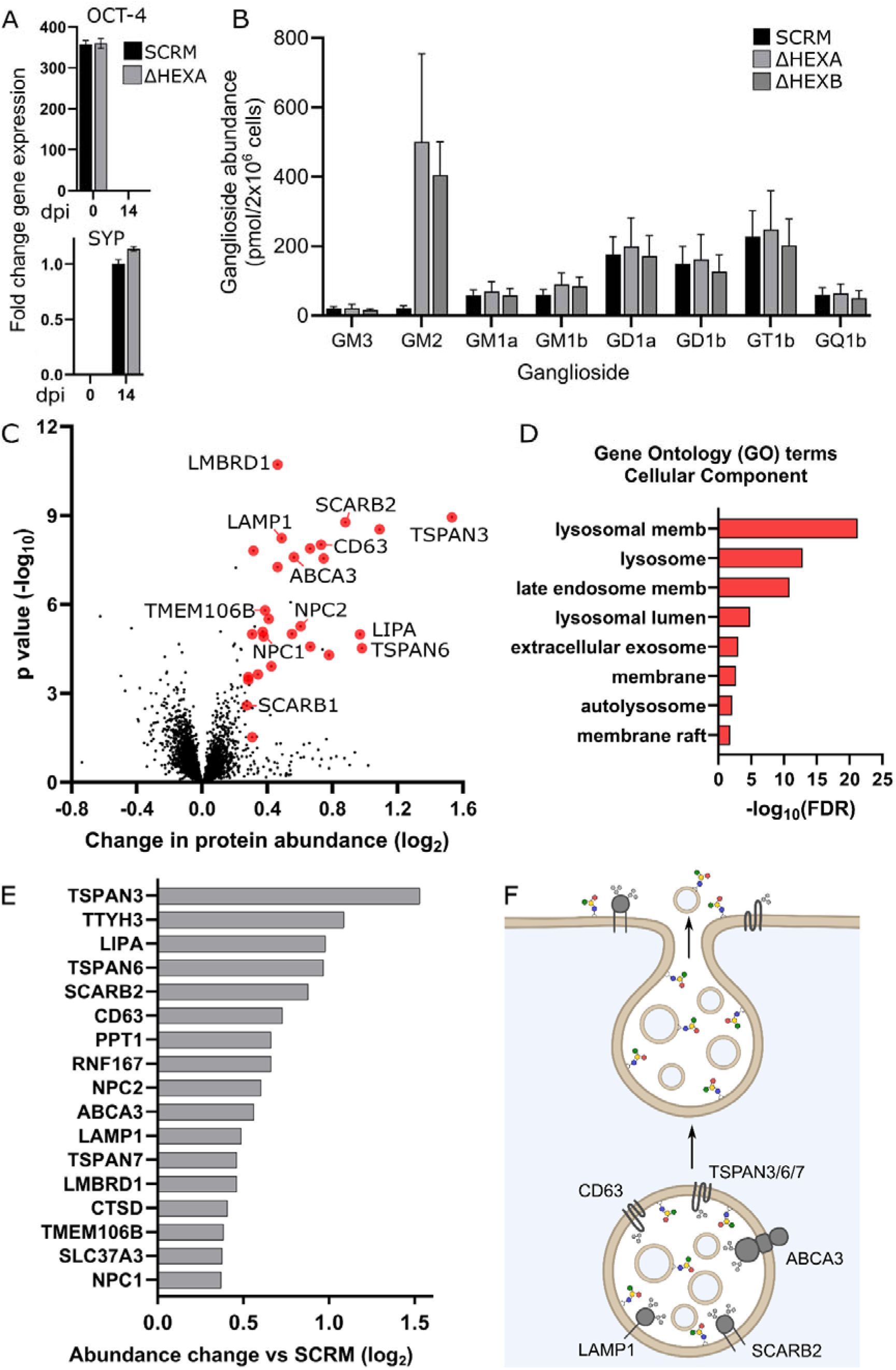
Neuronal maturation, GM2 quantification and the impact of GM2 accumulation on the whole cell proteome of Tay-Sachs and Sandhoff disease models. **A.** qPCR analysis of gene expression for the stem cell marker OCT-4 and the neuronal marker synaptophysin (SYP) in SCRM and ΔHEXA cell lines at 0 and 14 dpi. Fold change is calculated relative to 14 dpi SCRM controls, n=3 biological replicates were carried out in technical triplicate and the mean is displayed ± SEM. **B.** Quantification of whole-cell gangliosides at 14 dpi for SCRM, ΔHEXA and ΔHEXB cell lines, the mean of n=3 biological replicates is displayed ± SEM. **C.** Quantitative whole cell proteomics (WCP) data for ΔHEXA and ΔHEXB neurons compared with the SCRM control. A volcano plot is shown with average fold change (x-axis) across three biological replicates and significance (y-axis, two-sided t test) across the three replicates. Endolysosomal proteins are coloured in red with targets mentioned in the text labelled. **D.** Gene ontology (GO) term analysis for cellular component of proteins significantly changed in the WCP dataset. **E.** Select targets from the WCP are represented graphically to illustrate the fold change in whole cell protein abundance in ΔHEXA and ΔHEXB neurons vs SCRM neurons. **F.** Schematic diagram of lysosomal exocytosis and how this process can contribute to changes in lipid and protein abundance at the plasma membrane (PM). A selection of lysosomal proteins increased in abundance in the WCP of ΔHEXA and ΔHEXB neurons are illustrated.

The quantification of complex glycosylated sphingolipids in cells is challenging using mass spectrometry approaches due to several glycan moieties possessing the same molecular mass [35]. An alternative strategy to accurately quantify the abundance of the different ganglioside headgroups is known as glycan profiling [36,37]. This involves extraction of all GSLs from cell lysates, cleavage of the ceramide tails using endocerebrosidase (ceramide glycanase), fluorescent labelling of the glycan headgroup with 2-Anthranilic Acid and separation of the labelled glycans using HPLC for quantification of individual peaks against known standards (**Supp. Fig. S2**). To determine how much GM2 was accumulating and whether any other gangliosides were altered in these disease models, glycan profiling was employed to quantify the abundance of all ganglioside species (**Fig. 2B**). These data confirm substantial accumulation (> 20-fold) of GM2 in ΔHEXA and ΔHEXB neurons, compared to SCRM and demonstrates that no other gangliosides are significantly altered in their abundance by the accumulation of GM2.

These data from i3N disease models match closely to the observed phenotypes in diseased patient neurons [21]. Specifically, the accumulation of GM2 lipid substrate, particularly in the late endolysosomes, causing formation of membrane whorls and enlargement of this compartment. Therefore, this is a valid model for probing the molecular mechanisms driving disease pathology.

### Impact on the proteome of GM2 accumulation in neurons

To understand the impact that GM2 accumulation has on neuronal function, whole cell proteomics (WCP) was carried out on SCRM, ΔHEXA-1/2 and ΔHEXB-1/2 at 14 dpi. High confidence targets were selected based on a significant change in abundance of at least 1.2-fold compared to SCRM control. When compared with each other, the models of Tay-Sachs (ΔHEXA-1/2) and Sandhoff (ΔHEXB-1/2) disease cell lines showed very few significant differences in protein abundances, indicating that the changes observed, compared to the SCRM line, are due to GM2 accumulation specifically rather than the exact gene/subunit that is targeted (**Supp. Fig. S3**). Thus, for analysis, data from both the models of Tay-Sachs (ΔHEXA-1/2) and Sandhoff (ΔHEXB-1/2) were combined and compared to SCRM controls. All disease models showed significant accumulation of endolysosomal proteins including degradative enzymes (such as LIPA and CTSD), membrane pumps (SLC37A3 and LMBRD1), lipid transporters (NPC1/2, SCARB2 and ABCA3), integral limiting membrane proteins (LAMP1), modulators of lysosomal position (RNF167 and TMEM106B) and exosomal markers (TSPAN3/6/7 and CD63) (**Fig. 2C** and **Supp. Table S2**). These changes in protein abundance were primarily due to the accumulation of endolysosomes rather than gene expression changes as demonstrated by qPCR data showing no significant changes between control and knockdown lines for five proteins involved in the CLEAR response (Coordinated Lysosomal Expression And Regulation) that drives lysosomal biogenesis (**Supp. Fig. S4**) [38,39].

The proteomic changes seen in these cells have been driven by lipid accumulation and the subsequent downstream effects that this accumulation has upon lysosomal function and cellular trafficking. Several of the proteins identified here, such as the tetraspanins (TSPANs), have recently been shown to directly bind GSL headgroups [40] and several others such as LIPA, PPT1, ABCA3 and NPC1/2 are directly involved in lipid and sphingolipid processing [41–44]. This suggests that some of the proteins that are accumulating in these diseases are specifically products of lipid accumulation rather than a product of general lysosomal dysfunction. In further support of this, several lysosomal proteins including V-type ATPases (ATP6 family), mannose-6-phosphate receptor (M6PR) and biogenesis of lysosomal organelle complex subunits (BLOC1) are quantified in the WCP but are not increased in abundance.

Functional annotation and enrichment analysis using the Database for Annotation, Visualisation and Integrated Discovery (DAVID) identified several gene ontology (GO) terms that were highly enriched in this dataset including lysosomal, endosomal and membrane terms (**Fig. 2D**). Of particular interest in the WCP datasets was the increasing abundance of a subset of proteins involved in endolysosomal trafficking and exocytosis as well as exosome formation and release including the TSPANs 3,6 and 7, CD63 and ABCA3 (**Fig. 2E** and **2F**) [45–47]. The exocytosis of endolysosomal components has a normal role in membrane repair and secretion but is also a strategy used by the cell to attempt to clear accumulating cell debris [48]. The fusion of these compartments with the PM may be beneficial for the cell and has been proposed as a mechanism that could be enhanced and exploited to treat LSDs [49]. However, the delivery of endolysosomal proteins and lipids to the PM may also result in undesired changes at the cell surface that may interfere with cellular function.

### Consequences of GM2 accumulation on the plasma membrane proteome

The increased abundance of cellular proteins involved in lysosomal trafficking and exocytosis may have significant consequences for the protein composition of the PM. The proteome of the PM makes up a small fraction of the total cellular proteome meaning that changes at the PM are often not quantifiable using whole-cell analytical techniques [50]. To ensure accurate quantification of the PM proteome, PM proteins were specifically labelled with aminooxy-biotin before cell lysis allowing for enrichment with streptavidin prior to mass spectrometry analysis. As before, all four cell lines, ΔHEXA 1/2 and ΔHEXB 1/2 were combined and compared against the control SCRM cell line at 14 dpi with high-confidence targets selected based on a significant change in abundance of at least 1.2-fold compared to SCRM control. Functional annotation of GO Terms was used to define cellular compartments and the data consisted overwhelmingly of membrane proteins, though some membrane adjacent proteins were also detected.

Significant increases in the abundance of several endolysosomal proteins was observed at the PM including proteins present in the limiting membrane including SCARB2, ACP2, NPC1 and TSPAN3 (**Fig. 3A, 3B** and **Supp. Table S3**). This provides strong evidence that endolysosomal compartments are fusing with the PM to disgorge undegradable lipid material, further supported by the increase in abundance of the lipid transporter CD36.

**Figure 3.**
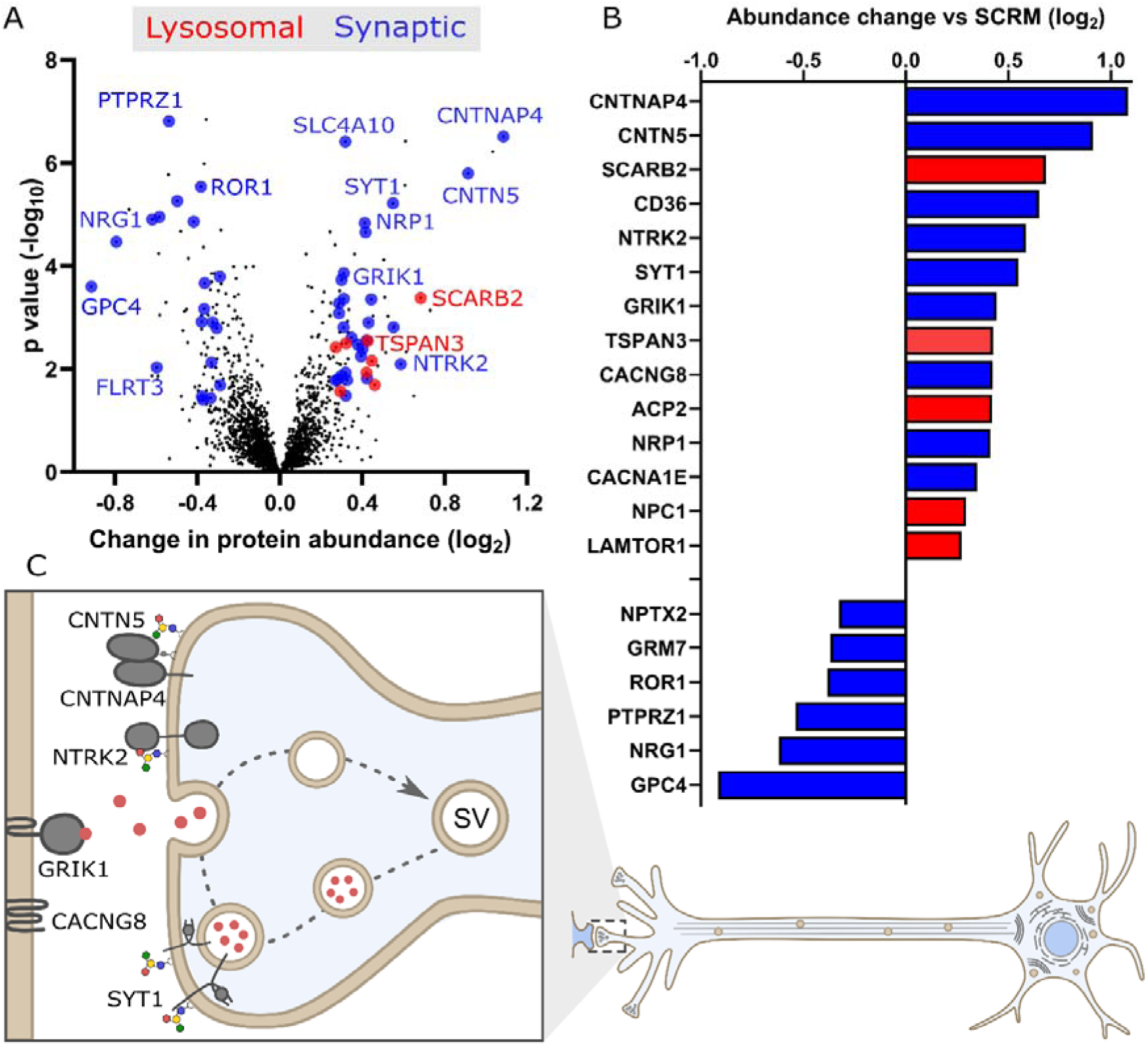
Molecular consequences of GM2 accumulation on the protein composition of the plasma membrane (PM). **A.** Quantitative mass spectrometry following enrichment of PM proteins from ΔHEXA and ΔHEXB neurons compared with the SCRM control. A volcano plot is shown with average fold change (x-axis) across three biological replicates and significance (y-axis, two-sided t test) across the three replicates. Targets coloured in blue are synaptic proteins and in red are lysosomal proteins with selected targets labelled. **B.** Select targets are represented graphically to illustrate the fold change in PM protein abundance in ΔHEXA and ΔHEXB neurons vs SCRM neurons. **C.** Illustration of proteins with increased abundance at the PM in ΔHEXA and ΔHEXB neurons, compared to SCRM neurons, that play important roles in neuronal signalling such as synaptic vesicle recycling and synaptic adhesion and receptor molecules.

Importantly, functional classification of the PM proteome data also revealed substantial additional changes at the PM including altered abundance of several synaptic proteins including “lipid raft”-associated and GPI-anchored proteins, such as CNTN5, CNTNAP4 (also known as Caspr4), IgSF21 and several glypicans GPC4/5/6 (**Fig. 3B** and **3C**). The protein abundance changes observed here at the PM are primarily driven by mislocalisation of proteins rather than altered gene expression as demonstrated by qPCR data for several high confidence targets (**Supp. Fig. S5**). Accumulating GM2 at the PM may be altering the formation or stability of membrane microdomains [51]. These membrane domains are critical for synaptic function including the clustering of synaptic proteins and typically contain large amounts of complex gangliosides that often interact directly with synaptic proteins [3,52,53]. Even small changes to the relative abundances of GM2 in these membrane microdomains may have a large impact on synaptic signalling and thus neuronal function.

In addition to proteins known to associate with membrane microdomains generally, some of the proteins identified here are direct ganglioside-binding proteins. The receptor tyrosine kinases NTRK2/3 (also known as TRKB/C) are significantly increased in abundance at the PM. NTRK2/3 promote neurite outgrowth, axonogenesis and synaptogenesis and also directly associate with gangliosides at the PM, typically GM1 [54]. NTRK2/3 are receptors for brain-derived neurotrophic factor (BDNF) and neurotrophin-3 [55] respectively and the increased abundance of these proteins at the PM in these disease models may be driving synaptogenesis and modulating neuronal activity.

Calcium flux across the PM and organelle membranes is known to be disrupted in both GM2 and GM1 gangliosidoses. Calcium sequestration is defective in both the ER and the lysosome in these disorders [56–59]. Several membrane proteins important in calcium signalling pathways are increased in abundance at the PM in these models of GM2 gangliosidoses including CACNA1E, CACNG8 and synaptotagmin 1 (SYT1). CACNA1E is a subunit of the 2.3 voltage gated calcium channel, driving calcium influx into cells at the pre- synapse to facilitate synaptic vesicle (SV) release, whilst CACNG8 positively regulates postsynaptic AMPAR activity [60–62]. The protein SYT1 is a calcium sensor required for neurotransmitter release and SV recycling at the synapse [63,64]. SYT1 and the SYT1- binding protein synaptic vesicle glycoprotein 2b (SV2b) [65,66] are both significantly increased in abundance at the PM in HEXA deficient cells. Previously, increased abundance of SYT1 at synapses was shown to drive faster SV internalisation and an increased SV exocytosis [67]. Similarly, the increase in abundance of SYT1 in these models of Tay-Sachs and Sandhoff diseases is likely to significantly impact synaptic signalling. Interestingly, SYT1 also specifically binds to the ganglioside GT1b suggesting there may be a direct link between ganglioside dysregulation and SV cycling [68].

Additional evidence of changes at the PM that are driven by altered GM2 metabolism and that may influence synaptic signalling in these neurons is the increased abundance of KCNA2, SCN2b and GRIK1 which are voltage and neurotransmitter gated potassium and sodium channels [69–71]. These increased abundances at the PM may have direct impact on the electrical conductance of the disease neurons.

### Molecular changes at the PM drive hyperactivity of Tay-Sachs neurons

To probe the impact of the PM proteome changes on neuronal signalling, initial measurements were made using a fluorescent calcium sensor protein [72]. Interestingly, this revealed that whilst SCRM, ΔHEXA-1 and ΔHEXB-1 neurons are beginning to fire spontaneously by 14 dpi, they do not undergo spontaneous synchronised signalling until 21- 28 dpi (**Fig. 4A**). This suggested that later time points might be more relevant for the study of mature synapse formation. Thus, although the neurons express markers of maturity by 14 dpi, it is not until 28 dpi that the neurons are functioning as a network. Over this longer time period, it is likely that GM2 accumulates to even greater quantities and the lysosomal exocytosis observed by 14 dpi suggests that this accumulation may also manifest at the PM. To probe alterations of the lipid composition of the PM, the ganglioside-profiling technique used previously (**Fig. 2B**) was modified to include an initial labelling step prior to cell lysis to distinguish cell-surface GSLs from intracellular GSLs. Using a modification of the technique used for labelling PM proteins, the sialic acid moieties of GSLs were labelled with an aminooxy-biotin that altered the elution profile of the cleaved headgroups during HPLC-based separation (**Supp. Fig. S6A**). Initial attempts to enrich for the PM fraction of lipids using streptavidin affinity were not successful meaning that the full repertoire of GSL content at the PM could not be determined. However, the quantification of several GSLs, including GM2, was possible as the aminoxy-biotin-labelled GSLs eluted in a region of the chromatogram free from other GSL peaks (**Supp. Fig. S6B**). Interestingly, this analysis demonstrated significant accumulation of GM2 at the cell surface of ΔHEXA cell lines, while other quantified gangliosides remained unchanged (**Fig. 4B**). By 14 dpi, the abundance of GM2 at the PM in ΔHEXA-1 cells is twice that in the SCRM cells (81 vs 38 pmol/2x10^6^ cells) and by 28 dpi increases to 5-fold higher than SCRM cells (248 vs 59 pmol/2x10^6^ cells). GM2 is not normally present in high amounts at the PM of mature neurons, and this increase in ΔHEXA-1 cells by 28 dpi brings its abundance up to the level of more common neuronal gangliosides such as GT1b.

**Figure 4.**
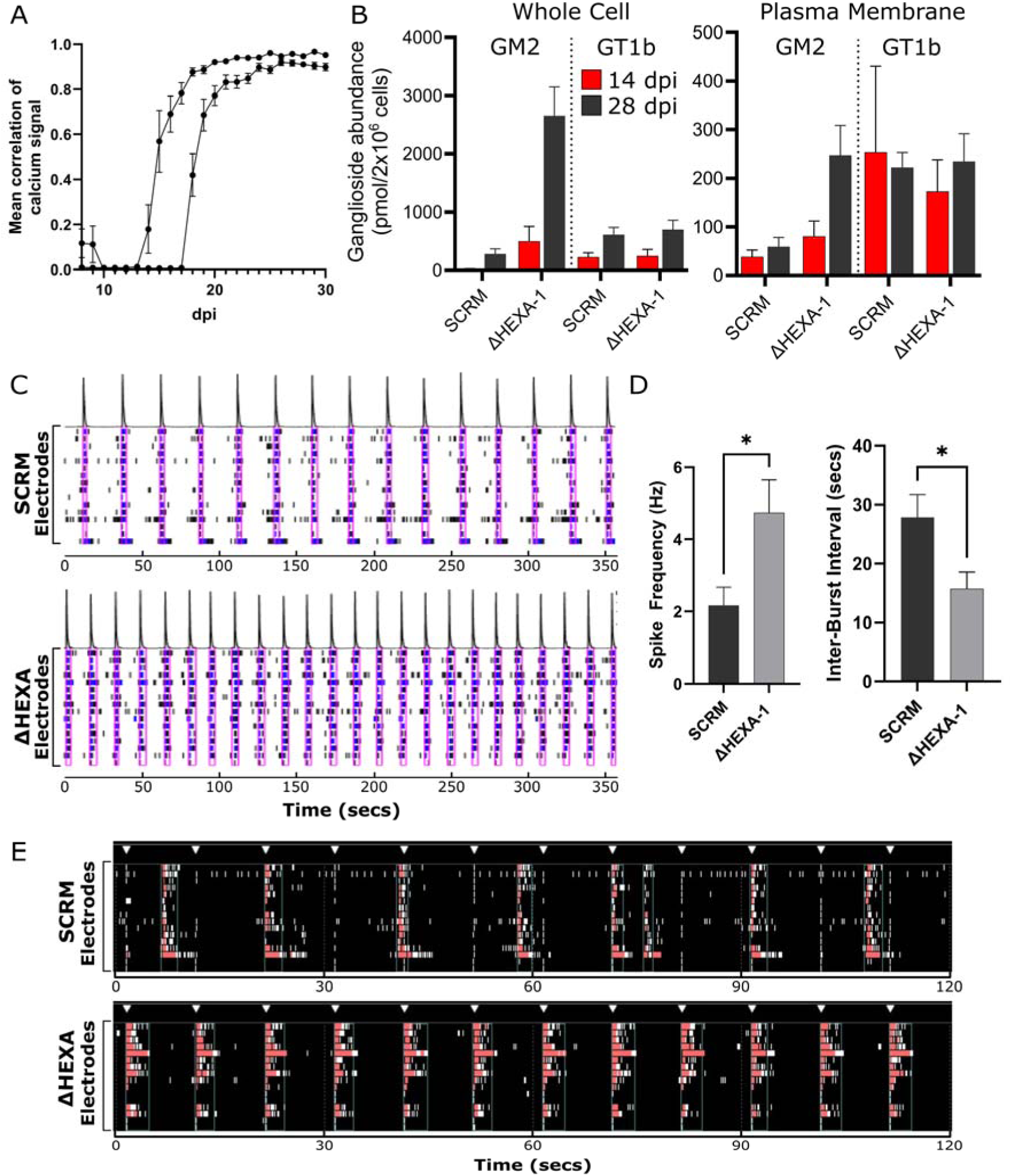
Increased spontaneous and stimulated neuronal network activity in. Δ**HEXA neurons compared with SCRM neurons. A.** Calcium signalling was monitored over 30 dpi for SCRM, ΔHEXA and ΔHEXB cell lines. Calcium signalling of cells starts to become synchronous (measured as correlation of bursting objects) from about 15 dpi with increasing signal correlation over time. Two independent experiments are shown with data from all cell lines (mean of SCRM, ΔHEXA and ΔHEXB ± SEM) combined to demonstrate that within an experiment, all cell lines are well correlated but between experiments, time to 100% correlation can differ until about 25 dpi. **B.** Quantification of gangliosides GM2 and GT1b from whole cell (left) and PM-enriched (right) samples of neurons at 14 and 28 dpi for SCRM and ΔHEXA cells. **C.** Representative activity traces (raster plots) of spontaneous neural activity for SCRM (upper) and ΔHEXA (lower) neurons. Activity is recorded over time on each of the fifteen electrodes with spikes (black), bursts (blue) and network bursts (pink boxes) shown over the same 360 second time period. The intensity of network activity is represented as a peak (above). **D.** Analysis of neuronal activity including spike frequency and inter-burst intervals were calculated covering 6 minutes of data collection. Data were collected from four wells of neurons per cell line and the mean is displayed ± SEM. **E.** Representative activity traces (raster plots) for SCRM (upper) and ΔHEXA (lower) neurons in response to repetitive field stimulation (white triangles). Activity over time is recorded on each of the fifteen electrodes. Network bursts (red) are shown over the same 120 second time period.

To explore the impact of changes in GM2 abundance and altered synaptic protein composition of the PM on neuronal activity, multi-electrode array (MEA) experiments were carried out on ΔHEXA-1 neurons. MEA allows for the monitoring of both spontaneous signalling of neurons and response to electrical stimulation. Following optimisation of the coating strategy and cell seeding, neurons were cultured on 24-well MEA plates to record neuronal activity. Electrical signalling was monitored via an array of electrodes with each electrode measuring spikes of field potential, and when > 5 spikes were measured on a single electrode within 100 ms these were considered bursts. When > 70% of electrodes in a well registered bursts within 100 ms this was considered a network burst. Consistent with the calcium signalling measurements, neurons were electrically active (bursting) by 14 dpi and firing synchronously (network bursting) by 21 dpi, with this burst strength increasing over time. At 42 dpi, ΔHEXA-1 neurons had significantly more frequent network bursts, firing faster than the SCRM control cells (**Fig. 4C** and **D**). Representative activity traces (raster plots) demonstrate the increased firing rate in ΔHEXA-1 cells and quantification over four wells of each cell line reveal significant differences in the mean spike frequency as well as inter-burst interval (mean interval dropping from 28 secs to 16 secs, *p* < 0.05).

The increased spontaneous burst rate for the diseased neurons is suggestive of a reduced threshold for electrical firing. To test whether this reduced threshold was recapitulated with field stimulation, neurons were subjected to electrical stimulation of 150 mV for 200 μs at a rate of 0.1 Hz for 5 minutes. The ΔHEXA-1 neurons again responded differently to field stimulation when compared with SCRM neurons. Control neurons did not respond readily to stimulation and continued to fire spontaneously at the same rate as they had prior to the applied stimulation (**Fig. 4E**, white triangles denote applied stimulation). In contrast, the ΔHEXA-1 cells consistently depolarise in synchronicity with the stimulation throughout the entire time course (**Fig. 4E**). The electrical stimulation was increased to 250 mV for 400 μs with the cell lines maintaining the same behaviour, that is ΔHEXA-1 neurons depolarising with each stimulation and SCRM neurons continuing to fire as they had spontaneously without stimulation. These data support that the ΔHEXA-1 neurons have a reduced threshold for depolarisation compared to SCRM cells. This hyperactive electrical signalling for the ΔHEXA-1 neurons is consistent with the increased abundance at the PM of key synaptic signalling proteins.

## DISCUSSION

The data presented here demonstrate the utility of the i3N cells for modelling neuronal diseases. The ΔHEXA and ΔHEXB neurons faithfully recapitulate the key disease phenotypes and are robust and reproducible enough to be used at scale to produce proteomic datasets to probe the mechanisms of GM2 gangliosidosis disease pathogenesis. The data establishes that whilst the lysosome is clearly the primary affected organelle in the GM2 gangliosidoses, the impact of lysosomal accumulation of lipids spreads beyond this organelle, with important implications for cellular and synaptic signalling in neurons. The changes observed at the PM are in part triggered by lysosomal dysfunction and impact both the lipid and protein composition of this membrane in complex ways. One of the major consequences of these molecular changes is altered synaptic signalling in disease neurons with them exhibiting hyperactivity due to a lower threshold for depolarisation.

The immunofluorescence and electron microscopy imaging for the HEXA-deficient cells revealed significant accumulation and enlargement of endolysosomal compartments. Interestingly, the more detailed analysis of these changes via WCPs revealed specific enrichment of lysosomal proteins involved in lipid processing and transport, including several proteins that bind directly to cholesterol and/or glycosphingolipids. This specific accumulation of proteins such as SCARB2, LIPA and NPC1/2 may represent “secondary” changes in response to GM2 accumulation. This suggests that rather than a general blockade of all lysosomal function, there may be a specific stabilisation, or reduced turnover, of lipid processing and lipid transporting proteins in an attempt to restore lipid homeostasis. The continued accumulation of GM2 and consequential impact on endolysosomal function also appears to trigger lysosomal exocytosis and exosome release as evidenced by increased abundance of trafficking and exosomal proteins in the WCP data and appearance of lysosomal proteins in the PMP datasets. These processes are mechanisms by which the cell attempts to relieve the lysosomal burden of accumulating substrate into the extracellular space [49]. Lysosomal exocytosis involves the wholesale fusion of the lysosomal limiting membrane with the PM and release of the contents, including the intra-lysosomal vesicles (ILVs) and enzymes, whilst exosome release involves the fusion of the multi-vesicular body (MVB) with the PM, releasing small (∼50-150nM diameter) vesicles into the extracellular space [73,74]. It seems likely that both are occurring in these disease models demonstrating that the consequences of lysosomal blockade are not confined to these endolysosomal organelles.

Two proteins identified in the WCP stand out as potentially very important in managing the lipid burden in these disease models: ABCA3 and the tetraspanin family member CD63. Both proteins are organised into membrane microdomains in the limiting membrane of endolysosomal compartments [75,76]. The lipid transporter ABCA3 is highly expressed in the brain but is well studied for its role in producing surfactant in the lungs [77–79]. Surfactant is a secreted lipid and protein mixture that coats the alveoli. Before secretion, surfactant components are packaged into lamellar bodies, a type of lysosome-related organelle (LRO) or specialised form of the MVB [80]. ABCA3 pumps lipids and cholesterol into lamellar bodies producing characteristic lipid whorls that are morphologically similar to those seen here (**Fig. 1F**) and in patient and animal models of sphingolipidoses. Importantly, ABCA3 can transfer a broad variety of lipids including phospholipids, cholesterol and the sphingolipids sphingomyelin and glucosylceramide [41]. This suggests that the increased abundance of ABCA3 observed in the HEXA-deficient cell lines may contribute to the formation of the multilamellar membrane structures seen in these cells. In addition to this packaging role, ABCA3 regulates the release of lamellar bodies via fusion with the PM: deletion of ABCA3 drastically reduces exosome release, whilst overexpression enhances release [45]. Therefore, it is possible that the increased abundance of ABCA3 observed in these diseased neurons is contributing to both lipid whorl formation and transport of GM2 between the endolysosome and PM. Interestingly, SCARB2, NPC1 and NPC2 are also found on lamellar bodies and function to translocate cholesterol and sphingolipids between endolysosomes and other membranous compartments, particularly the ER [81–84]. Thus, formation of lamellar-like bodies, similar to those in lung cells and seen in NPC deficiency [85], may be a mechanism by which storage and subsequent release occurs in the GM2 gangliosidoses.

Four tetraspanin proteins (TSPAN3/6/7 and CD63) were significantly upregulated in the WCP of HEXA deficient neurons. These are important scaffolding proteins for sorting and loading cargoes in MVBs and are enriched on exosomes and play important roles in exosome formation [46,47,86,87]. CD63 sorts cholesterol into ILVs to be used as a storage molecule or to be released via exosomes. NPC1/2 can then retrieve this cholesterol in either the cell-of-origin or after the exosome has been taken up by another cell [88]. The packaging and release of cholesterol in this manner is thought to be a mechanism by which exosomes are used to remove excess cholesterol in the LSD Neimann Pick disease type C (NPC) where cholesterol release in exosomes is increased [88]. Although CD63 is the best studied of these TSPANs, recently TSPAN3 was shown to be more efficient at sorting extracellular vesicles than CD63 [47]. Thus, the increased abundance of these tetraspanin proteins, seen in these neuronal disease models supports a significant role for exosome release in GM2 sphingolipidoses.

The proteomic profile of these cells paints a picture of specific dysfunction of lipid processing and an activation of alternative pathways to evacuate pathological GM2 to compensate for the lack of functional HexA. The relevance of these mechanisms to disgorge excess GM2 is supported by the specific and significant increase in abundance of GM2 seen in the PM glycan profiling. This showed continuing accumulation from 14 dpi increasing three-fold by 28 dpi as the cells exocytose their defective endomembrane compartments. This increase in GM2 at the PM is likely to be playing a specific role in disruption of membrane microdomains further contributing to the altered PM protein composition. There are several PM proteins that are altered in their abundance, both increased and decreased, that are not lysosomal proteins and so have been disrupted in their trafficking via pathways other than lysosomal exocytosis. These changes are likely driven by altered endo- and exocytosis of PM components due to the crucial role GSLs play in these recycling pathways [6,89]. In support of defective trafficking altering the PM proteome, several synaptic proteins (NTKR2, CACNG8, CNTNAP4, NRP1) were quantified in the WCP as not significantly changed but were changed in abundance at the PM, supporting a trafficking blockade with consequences for synaptic function.

Interestingly, the trafficking defects and therefore the proteomic changes at the PM may be specific to the type of ganglioside that is accumulating in these cells. Direct interactions between membrane proteins and ganglioside headgroups are difficult to define but there is substantial growing evidence that these interactions are specific and may help explain the different phenotypes seen in different GSL-related diseases [90]. Recently, the human ganglioside interactome for GM3, GM1 and GD1a was explored using photo-clickable GSLs [40]. This work identified membrane proteins that specifically bind to the different ganglioside-headgroups of these lipids. Interestingly, many of the proteins identified in our proteomics data including lysosomal (TSPAN3, CD63 and LAMP1) and non-lysosomal proteins (TTYH3, NTRK2/3, ROR1, EGFR amongst others) are direct ganglioside binders. GM2 is a ganglioside precursor that is not normally present at high abundance in the neuronal PM, thus the abundance increases seen here may disrupt the formation of correct ganglioside-protein interactions. The overlapping list of proteins identified here suggests that the excess GM2 may be disrupting normal GSL-protein interactions. The excessive amounts of inappropriately localised GM2 could compete with other GSLs to form non-native complexes that disrupt normal trafficking. Several ganglioside binding proteins have been shown to bind a range of gangliosides with different affinities and in this way interaction partners identified for GM3, GM1 and GD1a may be “hijacked” by excess GM2 [68,91].

The abundance changes at the PM for synaptic proteins involved in calcium signalling, synaptic vesicle recycling and neurotransmitter binding have clear consequences on the electrical activity of disease neurons. The ΔHEXA-1 neurons undergo more rapid network bursting both spontaneously and with extrinsic electrical stimulation. Synaptic alterations are well documented in LSDs but the molecular mechanisms driving this remain unclear. The work presented here supports that rather than a general neuronal dysfunction, there are specific changes to the composition of the synapse that in this neuronal type, cortical glutamatergic neurons, results in hyperactivity. Similarly, in a neuronal model of GM1 gangliosidosis it was shown that the architecture of the synapse is changed with the increase in GM1 abundance at the PM leading to downstream effects such as increased synaptic calcium levels and increased NTRK2 activity and subsequent increases in synaptic spine density [92]. In the data here for GM2 gangliosidoses, NTRK2 was also identified, suggesting there will be some common pathways and proteins affected in these related diseases.

Several studies using different cellular and animal models of LSDs have identified synaptic signalling defects with a range of different consequences depending on the specific disease and system studied [93,94]. In several cases reduced signalling is observed caused by fewer synaptic vesicles, loss of key synaptic proteins and loss of dendritic spines. In a recent study developing iPSC models from Tay-Sachs patient fibroblasts, exocytotic activity was reduced in response to chemical stimuli [26]. However, other work from NPC mice, display increased synaptic transmission potentially driving chronic excitotoxicity and neurodegeneration [95,96]. Enhanced exocytosis has also been observed in a mouse model of mucolipidosis IV supporting a role for glutamate neurotoxicity and synaptic exhaustion in neuronal pathology [97]. iPSC neuronal lines derived from different NPC patients have been shown to possess a higher resting voltage than healthy neurons and are easier to depolarise [98]. NPC neurons accumulate not only cholesterol but also multiple sphingolipids including GM2 [99,100]. A similar change is seen in the ΔHEXA-1 neurons here, where cells are more readily depolarised with electrical stimulation. Disruptions in synaptic function are challenging to study, but the system used here clearly demonstrates that GM2 accumulation in these neurons alters synapse composition such that they fire more frequently both spontaneously and with stimulation.

Enhancement of lysosomal exocytosis has been explored as a method to ameliorate the effects of LSDs via chemical induction, using cyclodextrin, or induced TFEB expression [101,102]. This approach has been used to reduce the accumulation of cholesterol, GM2 and GM3 in NPC disease models and GM2 in GM2 gangliosidosis models [27,103,104]. Although enhancing lysosomal exocytosis could effectively reduce the accumulation of storage materials, the deleterious consequences of altered PM lipid and protein composition suggest caution should be used in implementing this approach [105]. Furthermore, in other neurodegenerative diseases, the release of exosomes is known to pathologically “seed” bystander cells by passing misfolded toxic amyloid β and tau proteins to nearby cells [106]. In GM2 gangliosidoses, exosomes may pass GM2 to nearby cells such as other neurons, astrocytes and microglia, that are also unable to degrade the GM2 ganglioside leading to its accumulation, potentially driving dysfunction in other cell types.

The data presented here provide deep molecular insights into the mechanisms driving GM2 gangliosidosis pathology. We define the specific changes occurring within the endolysosomal compartments, the impact this has on the PM composition and the subsequent consequences for electrical signalling within the neuronal network. The i3N cell lines developed here represent robust, reproducible and versatile models for the study of GM2 gangliosidoses enabling broad, unbiased analysis of protein and lipid composition of different cellular compartments. These models represent an excellent resource for future research including drug screening to alleviate or reverse lysosomal and synaptic phenotypes. Furthermore, these models can be expanded to explore the specific changes that occur in similar gangliosidoses and more distantly related LSDs. This work clearly identifies that gangliosidoses are not just lysosomal storage diseases but are also complex plasma membrane disorders that directly affect the electrical activity and synaptic composition of neurons.

## METHODS

### Cell line maintenance

Human CRISPRi-i3N stem cells [28] containing a stably integrated dead Cas9 (dCas9) were maintained at 37°C and 5% CO_2_ with either daily media changes of complete E8 medium or triweekly media changes with complete E8 Flex medium, on plates coated with Matrigel diluted 1:50 in DMEM/F-12 HEPES. Cells were split using either 0.5 mM EDTA to detach cells, where they were replated as colonies in E8 medium, or Accutase, where they were replated as individual cells with E8 medium plus 10 μM Rock Inhibitor Y-27632 (Tocris).

Differentiation of stem cells into neurons was done as previously described, with slight modifications [28]. Initial differentiation was induced following Accutase treatment and seeding of 6-8x10^6 stem cells into a 10 cm Matrigel-coated dish using induction media (IM), comprised of DMEM/F12 HEPES, supplemented with 1x N2 supplement, 1x NEAA,1x Glutamax and 2 μg/ml Doxycycline. During initial plating, the media was supplemented with 10 μM Rock Inhibitor and media was changed daily for three days.

Partially differentiated neurons were detached with Accutase and replated onto plates coated with 100 μg/ml poly-l-ornithine (PLO), in cortical neuron (CN) media comprised of Neurobasal Plus medium supplemented with 1x B27 supplement, 10 ng/ml NT-3 (Peprotech), 10 ng/ml BDNF (Peprotech) and 1 μg/ml laminin. During initial plating, media was supplemented with 1 μg/ml doxycycline. Media was half changed twice or thrice weekly until day 14 unless otherwise specified.

### Generation of CRISPR knockdown cell lines

Guide RNAs targeting the *HEXA* and *HEXB* genes were designed using the CRISPick server (Broad Institute, **Supp. Table S1**) to target either the start of or immediately prior to the first exon and complimentary overhanging primer dimers were cloned into the BpiI restriction site on the pKLV plasmid (gift from Evan Reid).

For lentiviral production, HEK293T cells were maintained in DMEM supplemented with 10% FBS at 37°C and 5% CO_2_ and were passaged twice weekly with Trypsin-EDTA. 2x10^6 cells were transfected using 8 μl of Trans-IT and 2 μg of a 3:2:1 ratio of CRISPRi pKLV vector:pCMVΔ8.91:pMD.G lentiviral packaging vectors in Optimem. The transfection mix was added to cells dropwise and DMEM was replaced with E8 media after 24 hours. After 48 hours, virus containing E8 media was sterile filtered, diluted 1:1 with Fresh E8 and was frozen at -70°C.

5x10^5 i3N stem cells were transduced with 2ml of 1:1 E8 media/lentivirus-containing media supplemented with 10 μM Rock Inhibitor and 10 μg/ml Polybrene and spun onto Matrigel-coated 6-well plates at 750 *g* for 1 hour at 32°C, then incubated at 37°C. After 24 hours, media was changed with fresh E8 media and then 48 hours post transduction, media was changed with E8 medium supplemented with 1 μg/ml puromycin, daily for two days. Selected cells were subsequently grown without puromycin.

### Quantitative PCR (qPCR)

RNA was extracted from 2x10^6 cells as per manufacturer’s instructions using a Purelink RNA extraction kit (Invitrogen). RNA was reverse transcribed to cDNA using a High- Capactity RNA-to-cDNA kit (Applied Biosystems, Thermo Scientific), as per manufacturer’s instructions. RT-qPCR was performed using TaqMan Fast Advanced Mastermix (Applied Biosystems, Thermo Scientific), 50ng cDNA and FAM-labelled Taqman Gene Expression Assay probes (Applied Biosystems, Thermo Scientific) specific to each gene (**Supp. Table S4**). DNA amplification was performed using a CFX96 Real Time System (Bio-Rad) using 40 cycles of melting at 95°C for 15 seconds, annealing and extension at 60°C for 1 minute. Experiments were conducted with technical and biological triplicates. Gene expression changes were calculated using the ΔΔCt method [107] and using GAPDH as control. Statistical significance of fold changes were determined using the one-way ANOVA statistical test in Graph-pad Prism 9.

### Enzyme activity assays

HEXA activity assays were modified from previously published assays [32] as follows. Cells were lysed in 10 mM Tris pH7.4, 1% Triton X-100, 150 mM NaCl, 5 mM EDTA, supplemented with 1 cOmplete protease inhibitor tablet (Sigma Aldrich) per 50ml buffer and were centrifuged (12,000 *g,* 10 minutes) to remove debris. Volumes of cleared lysates were normalised following Pierce BCA assay (Thermo Fischer) and diluted to 40 μg/ml in PBS with 0.5% BSA. Samples were divided in two, with one half heated to 51.5°C for one hour. 4- Methylumbelliferyl-2-Acetamido-2-Deoxy-β-D-Glucopyranoside (MUG, Sigma Aldrich) or 4- Methylumbelliferyl-6-sulfo-N-acetyl-β-D-glucosaminide (MUGS, Sigma Aldrich) substrates were dissolved in 0.2 M Na_2_HPO_4_, pH 4.4 to a concentration of 3 mM. Heated samples were added to MUG substrate and both heated and unheated samples were added to MUGS substrate at a ratio of 1:2 v:v sample:substrate ratio. Lysate/substrate mixes were incubated at 37°C for 30 minutes at 200RPM in the dark before quenching with a 1:12 (v:v) sample:stopping buffer (250mM glycine carbonate, pH10.0) ratio. Fluorescence was read on a Clariostar plate reader (BMG LabTech) with excitation and emission wavelengths of 360 and 450 nm respectively. Experiments were conducted with technical and biological triplicates and statistical significance of differences in activity levels were determined using the one-way ANOVA statistical test in Graph-pad Prism 9.

### Quantification of gangliosides by HPLC

Glycosphingolipids were analysed as previously described [37,108]. Briefly, lipids were extracted from samples of 2x10^6 cells using 4:8:3 Chloroform:Methanol:PBS and further purified using solid phase C18 columns (Telos, Kinesis). Fractions were eluted twice with 1 ml of 98:2 Chloroform:methanol, and twice with 1 ml of 1:3 Chloroform:Methanol and 1 ml Methanol, then dried down under a nitrogen stream and digested overnight using recombinant endoglycoceramidase I (a gift from David Priestman). Cleaved glycans were labelled with 2-Anthranilic Acid (2AA) and purified using a DPA-6S Solid Phase Extraction amide column (Supelco). The purified, 2AA-labelled glycans were then separated and quantified by normal-phase HPLC and areas under the curve for each peak were compared to 2.5 pmol of 2AA-labelled chitotriose standard (Ludger) for quantification. For comparison between samples, this quantification was corrected for the number of cells seeded. For glycan identification, peak elution times were compared to a pre-determined mix of commercially available gangliosides (**Supp. Fig. S2**). Separation was performed using a TSK gel-Amide 80 column (Anachem) in combination with a Waters Alliance 2695 separations module and in-line Waters 474 fluorescence detector, set to excitation and emission wavelengths of 360 and 425 nm, respectively. All chromatography was performed at 30°C and solvent composition, gradient conditions and run parameters are as described previously [108]. Experiments were performed in biological triplicate and significant differences in the quantity of each ganglioside species were determined using the one-way ANOVA statistical test in Graph-pad Prism 9.

### Quantification of plasma membrane glycosphingolipids

Biotinylation of PM gangliosides was performed by modification of methods described previously for labelling of cell surface glycosylated proteins [109]. Briefly, 2x10^6 cells were washed twice in ice-cold PBS. Surface sialic acid residues were oxidised and biotinylated by incubation in the dark for 30 minutes with gentle rocking at 4°C using an oxidation/biotinylation mix comprising 1 mM sodium meta-periodate, 100 µM aminooxy-biotin (Biotium Inc., Hayward, CA) and 10 µM aniline (Sigma-Aldrich) in ice-cold PBS, pH 6.7. The reaction was quenched with 1 mM glycerol and cells were washed twice in ice-cold PBS. Labelled ganglioside headgroups were then purified and quantified using methods as described above for whole-cell analysis. Attempts to enrich and separate biotinylated gangliosides or cleaved glycan headgroups using streptavidin enrichment were unsuccessful. However, due to the altered elution profile of biotinylated glycan headgroups specific subsets of gangliosides were able to be quantified within the lipid mixture (**Supp. Fig. S6**). Identification of specific biotinylated glycan headgroups was determined by in vitro labelling of pure lipid species in liposomes composed of 20% ganglioside, 58% phosphotidylcholine, 20% cholesterol and 2% rhodamine-PE (**Supp. Table S5**).

### Transmission electron microscopy

Cell monolayers were fixed in 50% cell culture medium and 50% EM fixative (2% (w/v) paraformaldehyde and 2.5% (v/v) glutaraldehyde in 100 mM sodium cacodylate buffer, pH 7.2 at 37°C and this medium was immediately replaced with warm 100% fixative for 1h. Cells were washed with 100 mM sodium cacodylate buffer and post-fixed with 1% OsO_4_ in 100 mM sodium cacodylate buffer, pH 7.2 for 1h at room temperature. They were then washed with sodium cacodylate buffer followed by 50 mM sodium maleate buffer (pH 5.2) and ‘en bloc’ stained with 0.5% (w/v) uranyl acetate in 50 mM sodium maleate buffer, pH 5.2. The cells were sequentially dehydrated using 50% through to 100% ethanol and embedded in 50:50 v:v ethanol:Agar 100 resin (Agar Scientific) and finally 100% Agar 100 resin for 24h. A resin-filled BEEM capsule was then inverted over the cell monolayer and the resin was polymerised in an embedding oven for 24h at 60°C. The BEEM capsules were removed by immersion in liquid nitrogen.

Ultrathin sections were cut ‘en face’ to the plane of the monolayer using a diamond knife mounted on a Leica Ultracut UC7 ultramicrotome (Leica, Milton Keynes, UK), transferred to EM grids and stained with uranyl acetate and Reynold’s lead citrate. The sections were analysed using a Tecnai G2 Spirit BioTWIN transmission electron microscope (FEI) at an operating voltage of 80 kV and images were recorded using a 4 Megapixel Gatan US1000 CCD camera.

20 images per cell line were captured at 13,000x magnification. Images were analysed with ImageJ software, using a scale bar to define size with the analyse>set measurements function. The freehand selection tool and analyse>measure functions were then used to quantify endolysosome numbers and determine 2D endolysosome area measurements. Data were analysed by one-way ANOVA using GraphPad Prism.

### Immunofluorescence microscopy

Partially differentiated 3 dpi neurons were seeded onto coverslips in 24-well plates. Prior to seeding, coverslips were acid etched with 100% acetic acid and tumbling for 24 hours, then washed with ethanol and coated with 100 μg/ml PLO for 24 hours. At 14 dpi, coverslips were fixed with Cytoskeletal Fixing Buffer (300 mM NaCl, 10 mM EDTA, 10 mM Glucose, 10 mM MgCl_2_, 20 mM PIPES, pH 6.8, 2% Sucrose, 4% paraformaldehyde) for 10 minutes at RT. Coverslips were permeabilised with 0.1% Saponin in PBS for 10 minutes followed by blocking in 0.01% saponin, 1% BSA in PBS, and stained in the same buffer, with specific antibodies described below. Coverslips were stained for 1 hour in primary antibody, followed by three washes in 0.01% saponin in PBS, following by staining with secondary antibody for 45 minutes. Coverslips were then washed three times in 0.01% saponin in PBS and mounted in Prolong Gold Antifade with DAPI (Thermo Fisher). Antibodies used were rabbit polyclonal anti-LAMP1 (Abcam, ab24170), mouse monoclonal anti-GM2 (gift from Kostantin Dobrenis), AF555 donkey anti-rabbit (Invitrogen, A31572) and AF488 goat anti-mouse IgM (Invitrogen, A21042).

All images were collected on a Zeiss LSM 780 laser scanning microscope using Zen Black software and a 63x oil immersion lens and 512 x 512 pixel size images in a 16-bit range were obtained. Automated analysis and object-based colocalisation of image sets was performed using CellProfiler software [110]. The methodology is based on the CellProfiler online tutorials (https://tutorials.cellprofiler.org/) and the Cell Profiler Workflow is included as a Supplementary Data File. Briefly, images were split into colour layers with cells identified on the blue layer as objects between 25-80 pixels in diameter. Objects with diameters of 2- 20 pixels on the red layer (LAMP1 positive) were identified as endo/lysosomes and on the green layer as GM2. Colocalization was determined by identifying which green objects overlay with red. Counts for Nuclei, GM2, LAMP1 and GM2 positive for LAMP1 were used on a per image basis to produce GM2/cell and GM2 in lysosomes/Cell variables.

### Calcium activity assay

Partially differentiated 3 dpi neurons were seeded in 100 μl of media into PLO-coated 96- well plates (Corning, 3595) at a density of 5x10^4/well. 24 hours after seeding, 100 μl of diluted (1:100 v:v in CN media) Neuroburst Orange Lentivirus (Sartorius) expressing a genetically-encoded fluorescent calcium sensor was added to each well. 24 hours after virus addition, all media was removed and replaced with 200 μl of fresh CN media. Wells were imaged daily in a Incucyte S3 Live-Cell Analysis System (Sartorius) using movie mode acquisition for 2 minutes/well with 3 images/second, 4x objective lens, 400 ms acquisition time until 30 dpi. Analysis of the recorded movies was performed using the Neuronal Activity Analysis Software Module [72]. Briefly, active objects (cells/cell clusters) that burst above a minimum threshold were compared to every other object in the image to generate a correlation value between -1 and 1 with 1 being highly synchronous indicating high network connectivity. The following analysis parameters were chosen: object size 30 μm; minimum cell width 6 μm; sensitivity -3; edge split off; minimum burst intensity 0.2. Correlation scores for technical duplicates of SCRM, ΔHEXA-1 and ΔHEXB-1 were combined on a per experiment basis and presented to identify the time points where cells were signalling in high synchronicity.

### Proteomics

Detailed methods for sample preparation, TMT labelling, mass spectrometry and analysis of proteomics data are available in the Supplementary Information.

### Multielectrode array measurements

Wells of a 24-well MEA plate (Axion Biosystems) were coated with 100 μl of 100 μg/ml PLO and plates were incubated at room temperature overnight. Wells were washed three times with sterile water before being left to dry for approximately 1 h. 4x10^5 neurons/well were seeded in 100 μl of CN Media /maintenance media +1ug/ml doxycycline and maintained as described above in 200 μl of CN media. A Maestro MEA EDGE system and the control software Navigator (Axion BioSystems) were used for signal recording and stimulation. Spike detection was performed with a threshold of 6 times standard deviations. Electrical activity of neurons was monitored regularly for activity and synchronicity. When recordings took place on the same day as the half-media changes, neurons were recorded prior to changing the media as changes temporarily alter neuronal activity. On the recording day, the plate was loaded into the MEA reader, equilibrated for 15 min at 37°C and 5% CO_2_. For spontaneous signalling measurements, recording was performed via AxIS for 15 min (200 Hz–3 kHz). To electrically stimulate the culture, biphasic voltage pulses defined in the Navigator software (‘Neural Stimulation’ setting, amplitude = 150-300 mV, duration = 200- 400 μS) were delivered to the target electrodes at 0.1 Hz for 5-10 minutes. Recorded data were processed with the Navigator software and then subsequently using GraphPad Prism 9 software.

## Acknowledgements

This work was supported by a Wellcome Trust Senior Research Fellowship (219447/Z/19/Z) to JED that also supports ASN, HGB and ES. F.M.P. is a Wellcome Trust Investigator in Science and a Royal Society Wolfson merit award holder. D.A.P. was funded by the Mizutani Foundation. For the purpose of open access, the author has applied a Creative Commons Attribution (CC BY) licence to any Author Accepted Manuscript version arising from this submission.

## Data Availability

The mass spectrometry proteomics data produced in this study will be deposited to the ProteomeXchange Consortium via the PRIDE repository (https://www.ebi.ac.uk/pride/) with the dataset identifiers TBA. Glycan analysis data will be supplied with this submission as supplementary data files. Raw electron microscopy images will be deposited in the University of Cambridge Data Repository (TBA).

