## Supplementary Information for "Plasma Membrane Remodelling in GM2 Gangliosidoses Drives Synaptic Dysfunction"

**This document contains:**

Supplementary Figures S1-S6

Supplementary Tables S1-S5

Supplementary Methods

**Additional Supplementary Data File:**

Cell Profiler Workflow for data in Fig 1

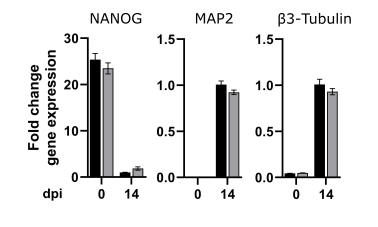

**Supplementary Figure S1.** qPCR analysis of gene expression for the stem cell marker NANOG and neuronal markers MAP2 and β3-tubulin in SCRM and ΔHEXA cell lines at 0 and 14 dpi. Fold change is calculated relative to 14 dpi SCRM controls, n=3 biological replicates were carried out in technical triplicate.

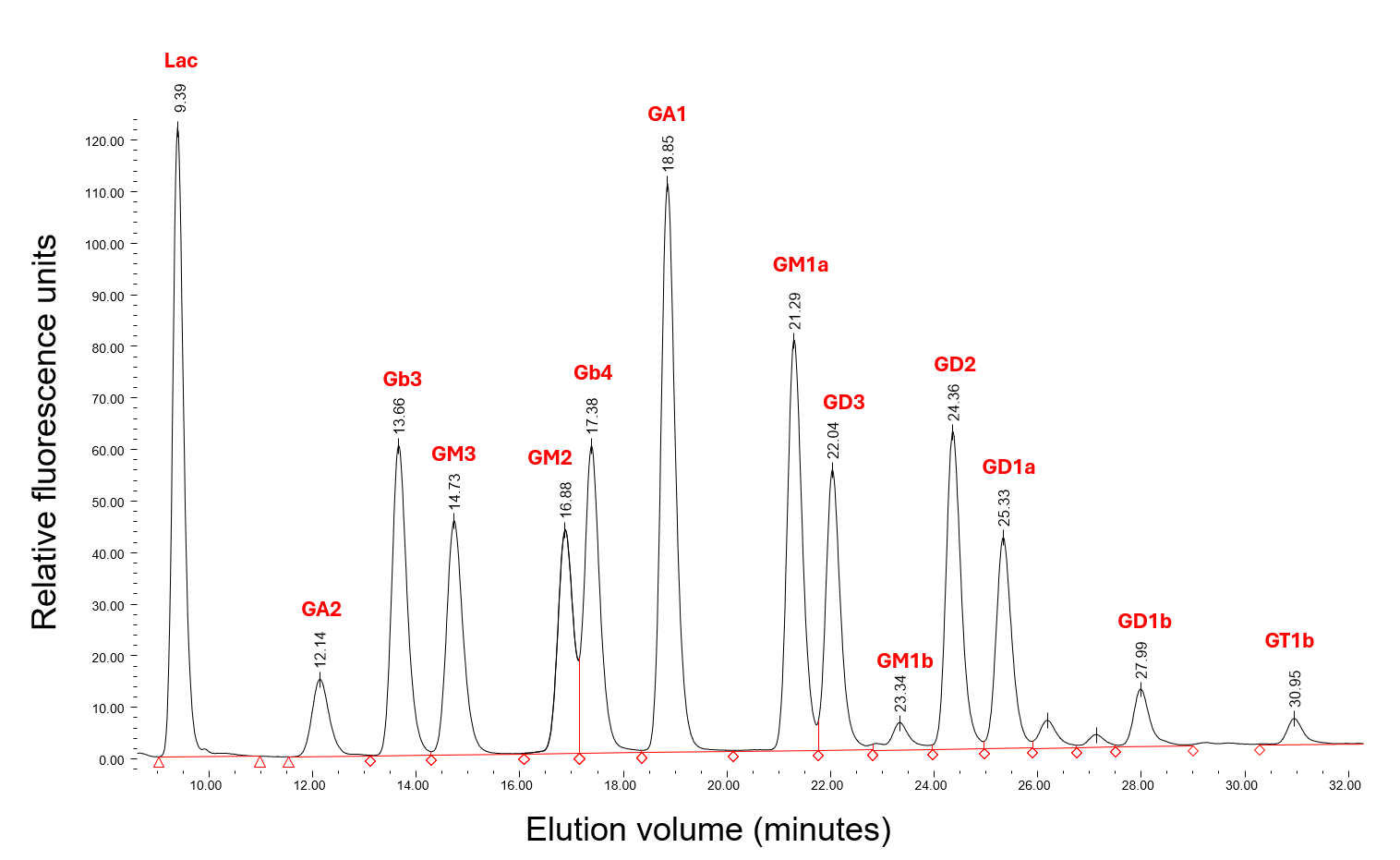

**Supplementary Figure S2.** Identification and quantification of glycosphingolipid headgroups using HPLC analysis of cleaved and labelled glycan headgroups. Elution profile of known GSL standards as reference for peak identification. Each glycan headgroup has a distinct elution volume (labelled).

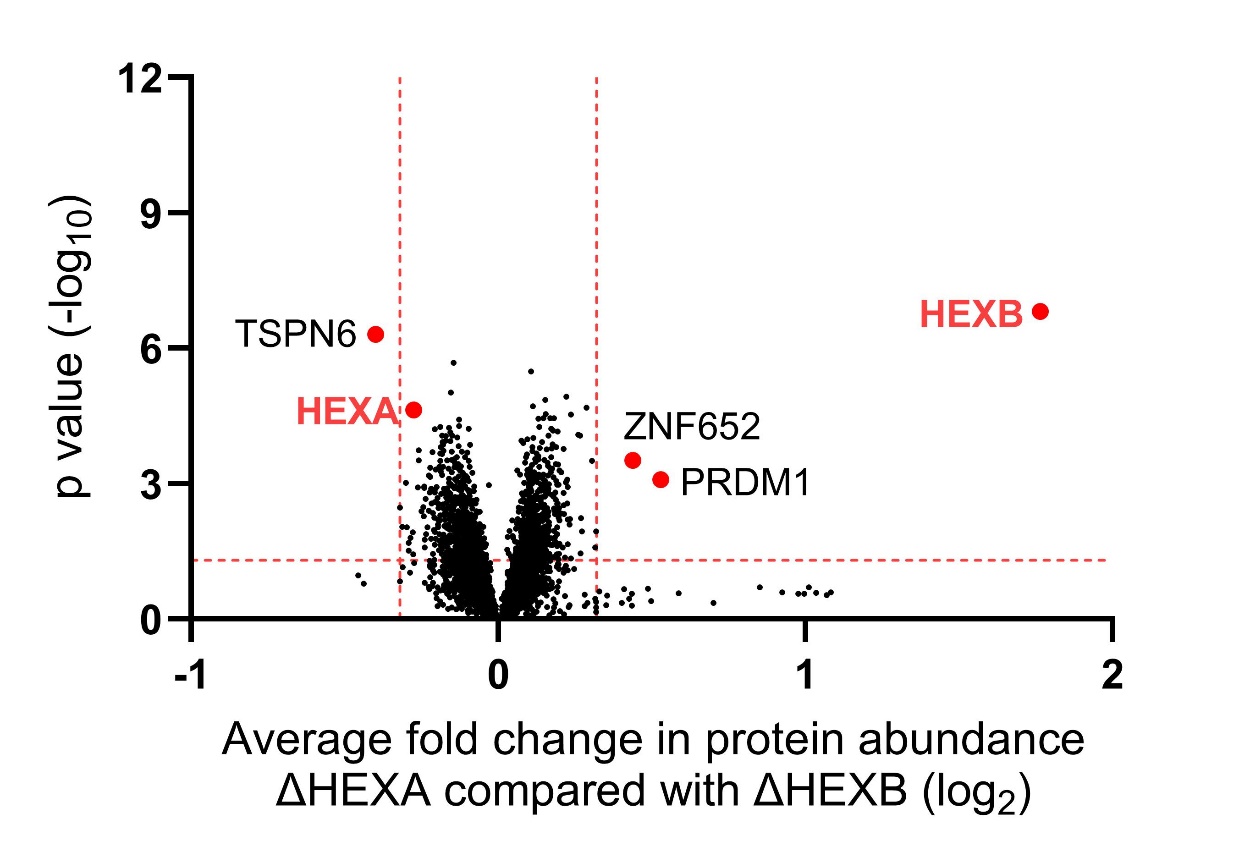

**Supplementary Figure S3.** Quantitative mass spectrometry from whole cell samples of ΔHEXA compared to ΔHEXB cell lines. A volcano plot is shown with the horizontal axis showing average fold change across three biological replicates and the vertical axis showing significance (two-sided t test) across the three replicates. Significance cutoffs of >0.25 fold change and p-value <0.05 are indicated (red dotted lines). Beyond the subunits that have been knocked down there are only 3 proteins with significant changes in protein abundance between these cell lines.

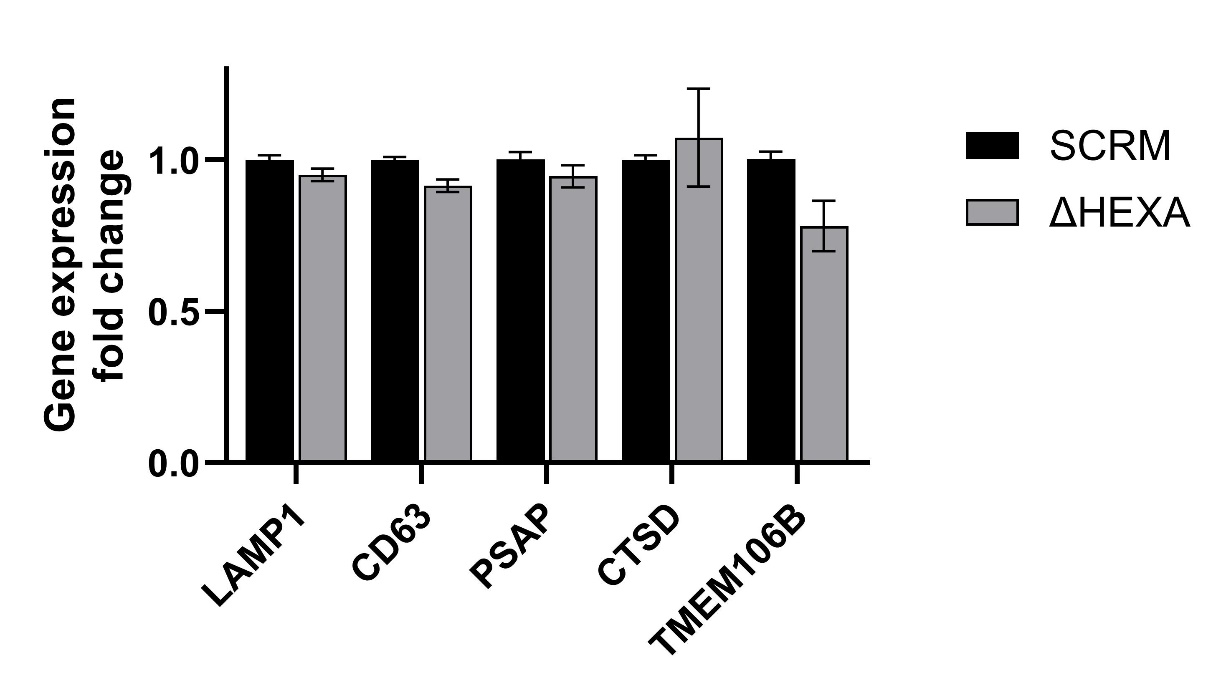

**Supplementary Figure S4.** qPCR analysis of a selection of lysosomal proteins involved in the CLEAR response including two that are increased in abundance in WCP data. Gene expression levels are shown at 14 dpi for SCRM and ΔHEXA lines. Fold change is calculated relative to SCRM controls, n=3 biological replicates were carried out in technical triplicate. No significant differences are seen between SCRM and ΔHEXA lines.

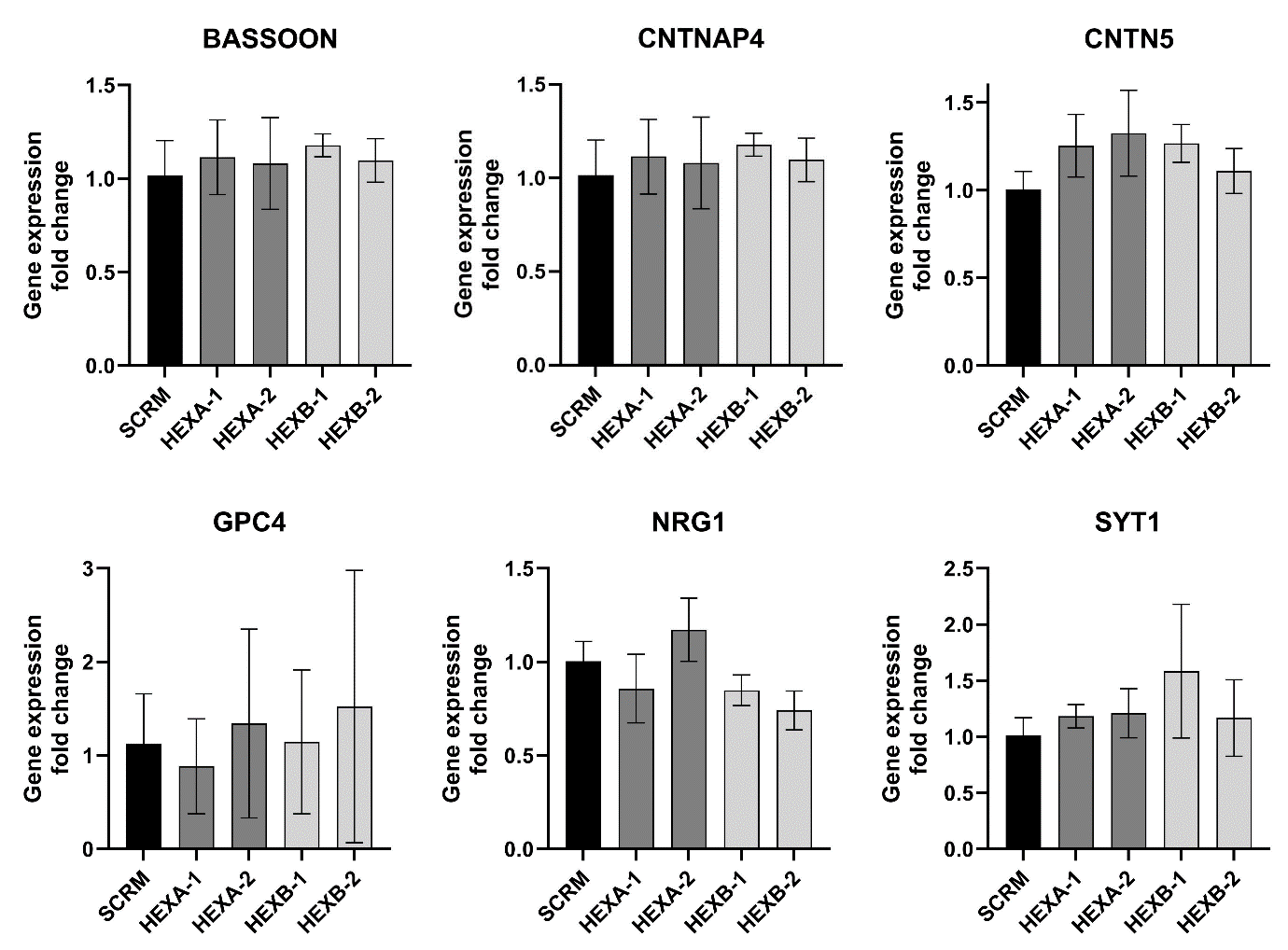

**Supplementary Figure S5.** qPCR analysis of a selection of high confidence targets identified as changed in abundance in PMP data at 14 dpi. n=3 biological replicates were carried out in technical triplicate.

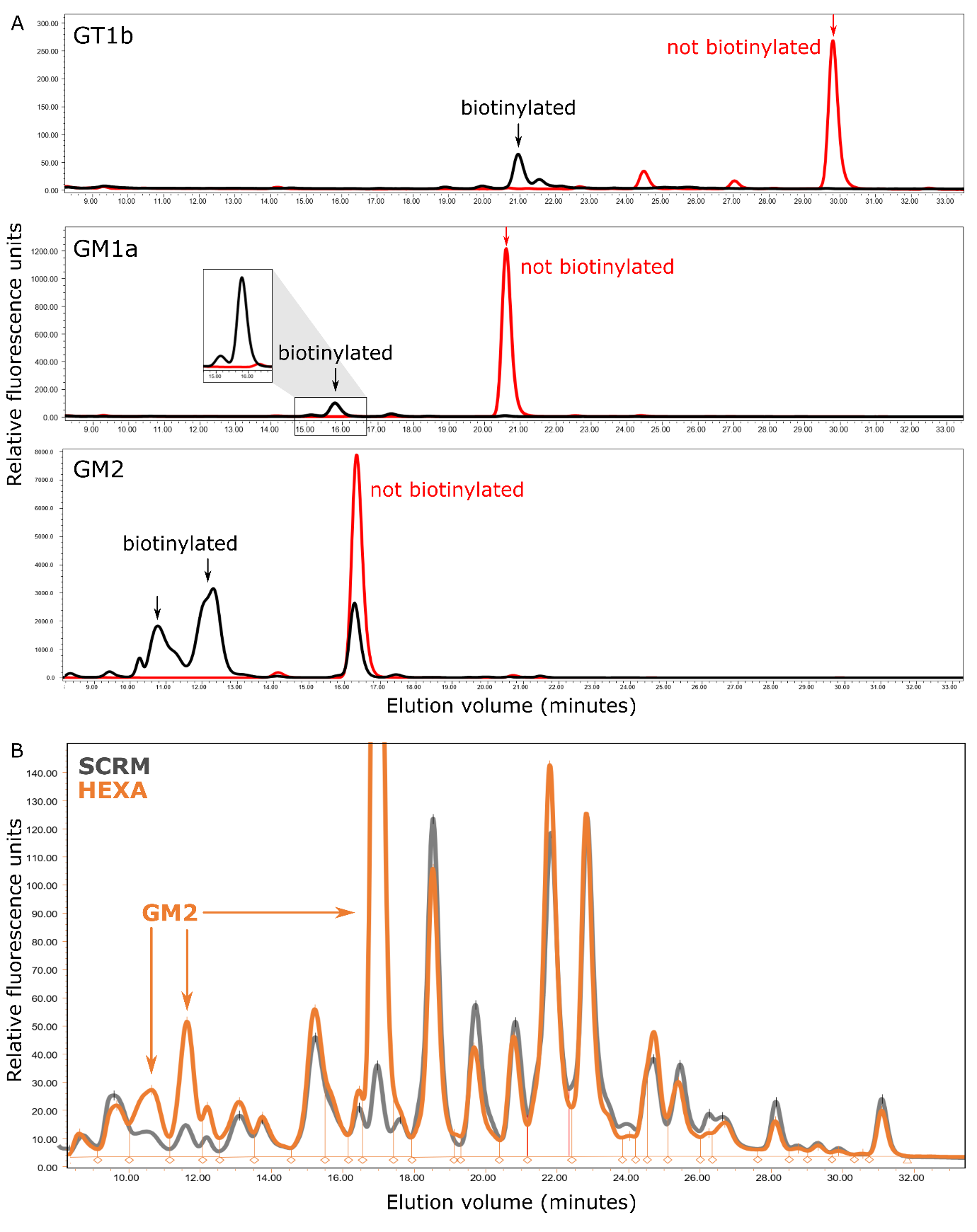

**Supplementary Figure S6.** HPLC elution profiles of biotinylated ganglioside headgroups. **A.** Standards from individually labelled lipid species GT1b, GM1a and GM2. **B.** Ganglioside headgroup elution profiles of 28 dpi whole cell samples after surface labelling with aminoxybiotin.

**Supplementary Table S1.** Guide RNA target sequences

| Gene | Exon | Target sequence |
| --- | --- | --- |
| HEXA-1 | 1 | 5’ - CAGGTCACGATAGCGCTGGA |
| HEXA-2 | 1 | 5’ - CCAAAGCCTGGAGCTTGTCA |
| HEXB-1 | 1 | 5’ - GCTGTTGGCGACACTGCTGG |
| HEXB-2 | Prior to 1 | 5’ - CCGCTCGGCTGCTTTCCGCC |

**Supplementary Table S2.** High confidence targets identified in WCP of ΔHEXA and ΔHEXB compared with SCRM control cells at 14 dpi.

| **Gene ID** | **Description** | **Log_2_ Fold change** | **Significance p-value** |
| --- | --- | --- | --- |
| O60637 | Tetraspanin-3 GN=TSPAN3 | 1.532 | 1.2E-09 |
| Q9C0H2 | Protein tweety homolog 3 GN=TTYH3 | 1.088 | 2.9E-09 |
| P38571 | Lysosomal acid lipase/cholesteryl ester hydrolase GN=LIPA | 0.980 | 3.0E-05 |
| O43657 | Tetraspanin-6 GN=TSPAN6 | 0.968 | 1.0E-05 |
| Q14108 | Lysosome membrane protein 2 GN=SCARB2 | 0.879 | 1.7E-09 |
| Q7Z3F1 | Integral membrane protein GPR155 GN=GPR155 | 0.778 | 5.1E-05 |
| P13645 | Keratin, type I cytoskeletal 10 GN=KRT10 | 0.763 | 3.5E-02 |
| Q9BXS4 | Trabsmembrane protein 59 GN=TMEM59 | 0.746 | 2.8E-08 |
| Q9BT67 | NEDD4 family-interacting protein 1 GN=NDFIP1 | 0.738 | 3.3E-05 |
| P08962 | CD63 antigen GN=CD63 | 0.730 | 9.6E-09 |
| P50897 | Palmitoyl-protein thioesterase 1 GN=PPT1 | 0.663 | 2.7E-05 |
| Q9H6Y7 | E3 ubiquitin-protein ligase RNF167 GN=RNF167 | 0.662 | 1.3E-08 |
| P31944 | Caspase-14 GN=CASP14 | 0.657 | 4.7E-02 |
| P61916 | Epididymal secretory protein E1 GN=NPC2 | 0.604 | 5.5E-06 |
| Q99758 | ATP-binding cassette sub-family A member 3 GN=ABCA3 | 0.563 | 2.5E-08 |
| O43567 | E3 ubiquitin-protein ligase RNF13 GN=RNF13 | 0.551 | 9.9E-06 |
| P78382 | CMP-sialic acid transporter GN=SLC35A1 | 0.543 | 8.3E-07 |
| P11279 | Lysosome-associated membrane glycoprotein 1 GN=LAMP1 | 0.488 | 5.8E-09 |
| Q96P63 | Serpin B12 GN=SERPINB12 | 0.481 | 4.9E-02 |
| P41732 | Tetraspanin-7 GN=TSPAN7 | 0.463 | 5.4E-08 |
| Q9NUN5 | Probable lysosomal cobalamin transporter GN=LMBRD1 | 0.462 | 1.9E-11 |
| Q8IY95 | Transmembrane protein 192 GN=TMEM192 | 0.425 | 1.2E-04 |
| P07339 | Cathepsin D GN=CTSD | 0.408 | 3.1E-06 |
| Q9NUM4 | Transmembrane protein 106B GN=TMEM106B | 0.386 | 1.6E-06 |
| Q8NCC5 | Sugar phosphate exchanger 3 GN=SLC37A3 | 0.377 | 1.2E-05 |
| O15118 | Niemann-Pick C1 protein GN=NPC1 | 0.371 | 8.4E-06 |
| P13473 | Lysosome-associated membrane glycoprotein 2 GN=LAMP2 | 0.342 | 2.3E-04 |
| P0CG05 | Ig lambda-2 chain C regions GN=IGLC2 | 0.337 | 2.9E-02 |
| Q9UJX6 | Anaphase-promoting complex subunit 2 GN=ANAPC2 | 0.323 | 5.5E-06 |
| Q96QD8 | Sodium-coupled neutral amino acid transporter 2 GN=SLC38A2 | 0.308 | 3.2E-03 |
| Q13286 | Battenin GN=CLN3 | 0.307 | 3.0E-02 |
| Q12999 | Tetraspanin-31 GN=TSPAN31 | 0.307 | 1.0E-05 |
| Q9UQM7 | Calcium/calmodulin-dependent protein kinase type II subunit alpha GN=CAMK2A | 0.298 | 2.7E-03 |
| O95772 | MLN64 N-terminal domain homolog GN=STARD3NL | 0.283 | 3.5E-04 |
| Q86WC4 | Osteopetrosis-associated transmembrane protein 1 GN=OSTM1 | 0.283 | 2.8E-04 |
| Q8WTV0 | Scavenger receptor class B member 1 GN=SCARB1 | 0.274 | 2.6E-03 |
| P61513 | 60S ribosomal protein L37a GN=RPL37A | -0.281 | 1.0E-03 |
| P04921 | Glycophorin-C GN=GYPC | -0.286 | 9.8E-03 |
| O75943 | Cell cycle checkpoint protein RAD17 GN=RAD17 | -0.297 | 1.0E-02 |
| Q15125 | 3-beta-hydroxysteroid-Delta(8),Delta(7)-isomerase GN=EBP | -0.299 | 3.5E-02 |
| F8WCM5 | Insulin, isoform 2 GN=INS-IGF2 | -0.302 | 1.8E-02 |
| Q96A83 | Collagen alpha-1(XXVI) chain GN=COL26A1 | -0.308 | 2.7E-03 |
| O15344 | E3 ubiquitin-protein ligase Midline-1 GN=MID1 | -0.313 | 9.9E-04 |
| Q9P2G3 | Kelch-like protein 14 GN=KLHL14 | -0.314 | 2.5E-02 |
| Q8WTS6 | Histone-lysine N-methyltransferase SETD7 GN=SETD7 | -0.321 | 2.0E-02 |
| Q96Q91 | Anion exchange protein 4 GN=SLC4A9 | -0.351 | 3.3E-03 |
| Q9Y2D9 | Zinc finger protein 652 GN=ZNF652 | -0.356 | 4.7E-02 |
| P29373 | Cellular retinoic acid-binding protein 2 GN=CRABP2 | -0.382 | 6.7E-04 |
| Q9NYJ7 | Delta-like protein 3 GN=DLL3 | -0.390 | 8.7E-03 |
| Q14831 | Metabotropic glutamate receptor 7 GN=GRM7 | -0.395 | 7.4E-03 |
| O60663 | LIM homeobox transcription factor 1-beta GN=LMX1B | -0.434 | 6.3E-06 |
| P41145 | Kappa-type opioid receptor GN=OPRK1 | -0.472 | 8.2E-04 |
| O14522 | Receptor-type tyrosine-protein phosphatase T GN=PTPRT | -0.498 | 2.6E-04 |
| Q13237 | cGMP-dependent protein kinase 2 GN=PRKG2 | -0.625 | 2.5E-06 |
| P06865 | Beta-hexosaminidase subunit alpha GN=HEXA | -1.045 | 5.2E-08 |

**Supplementary Table S3.** High confidence targets identified in PMP-MS of ΔHEXA and ΔHEXB compared with SCRM control cells at 14dpi.

| **Gene ID** | **Description** | **Log_2_ Fold change** | **Significance p-value** |
| --- | --- | --- | --- |
| Q9C0A0 | Contactin-associated protein-like 4 GN=CNTNAP4 | 1.084 | 3.0E-07 |
| P21583 | Kit ligand GN=KITLG | 1.032 | 6.0E-07 |
| O94779 | Contactin-5 GN=CNTN5 | 0.913 | 1.6E-06 |
| Q14982 | Opioid-binding protein/cell adhesion molecule GN=OPCML | 0.729 | 7.2E-04 |
| Q14108 | Lysosome membrane protein 2 GN=SCARB2 | 0.683 | 4.1E-04 |
| P16671 | Platelet glycoprotein 4 GN=CD36 | 0.649 | 3.3E-02 |
| O43194 | G-protein coupled receptor 39 GN=GPR39 | 0.612 | 4.6E-03 |
| P55283 | Cadherin-4 GN=CDH4 | 0.609 | 3.7E-07 |
| Q86SJ2 | Amphoterin-induced protein 2 GN=AMIGO2 | 0.606 | 2.7E-06 |
| Q16620 | BDNF/NT-3 growth factors receptor GN=NTRK2 | 0.585 | 8.0E-03 |
| O60320 | Protein FAM189A1 GN=FAM189A1 | 0.555 | 1.5E-03 |
| P78333 | Glypican-5 GN=GPC5 | 0.551 | 1.5E-03 |
| P21579 | Synaptotagmin-1 GN=SYT1 | 0.549 | 6.1E-06 |
| Q96ID5 | Immunoglobulin superfamily member 21 GN=IGSF21 | 0.524 | 9.3E-04 |
| Q9H461 | Frizzled-8 GN=FZD8 | 0.513 | 4.7E-03 |
| Q9H3S1 | Semaphorin-4A GN=SEMA4A | 0.508 | 4.1E-04 |
| P78382 | CMP-sialic acid transporter GN=SLC35A1 | 0.504 | 7.5E-05 |
| O00451 | GDNF family receptor alpha-2 GN=GFRA2 | 0.475 | 8.7E-03 |
| Q13286 | Battenin GN=CLN3 | 0.460 | 2.0E-02 |
| Q9UHI5 | Large neutral amino acids transporter SS 2 GN=SLC7A8 | 0.452 | 9.2E-04 |
| P38571 | Lysosomal acid lipase/cholesteryl ester hydrolase GN=LIPA | 0.446 | 6.9E-03 |
| P39086 | Glutamate receptor ionotropic, kainate 1 GN=GRIK1 | 0.443 | 4.4E-04 |
| O94933 | SLIT and NTRK-like protein 3 GN=SLITRK3 | 0.430 | 1.2E-03 |
| O60637 | Tetraspanin-3 GN=TSPAN3 | 0.426 | 2.8E-03 |
| Q8WXS5 | Voltage-dependent calcium channel gamma-8 subunit GN=CACNG8 | 0.422 | 1.5E-02 |
| P11117 | Lysosomal acid phosphatase GN=ACP2 | 0.420 | 1.2E-02 |
| Q16288 | NT-3 growth factor receptor GN=NTRK3 | 0.420 | 2.8E-03 |
| P28222 | 5-hydroxytryptamine receptor 1B GN=HTR1B | 0.415 | 2.2E-05 |
| O14786 | Neuropilin-1 GN=NRP1 | 0.412 | 1.5E-05 |
| Q9HCJ2 | Leucine-rich repeat-containing protein 4C GN=LRRC4C | 0.400 | 4.0E-03 |
| O43300 | Leucine-rich repeat transmembrane neuronal protein 2 GN=LRRTM2 | 0.393 | 5.6E-03 |
| P08913 | Alpha-2A adrenergic receptor GN=ADRA2A | 0.378 | 3.3E-03 |
| Q9Y6N7 | Roundabout homolog 1 GN=ROBO1 | 0.361 | 5.8E-05 |
| Q15878 | Voltage-dependent R-type calcium channel subunit alpha-1E GN=CACNA1E | 0.348 | 2.5E-03 |
| Q32ZL2 | Lipid phosphate phosphatase-related protein type 5 GN=LPPR5 | 0.335 | 7.0E-04 |
| Q6ZN44 | Netrin receptor UNC5A GN=UNC5A | 0.331 | 2.0E-03 |
| Q7Z4T9 | Protein MAATS1 GN=MAATS1 | 0.330 | 6.3E-03 |
| Q86TG7 | Retrotransposon-derived protein PEG10 GN=PEG10 | 0.328 | 1.2E-03 |
| O43295 | SLIT-ROBO Rho GTPase-activating protein 3 GN=SRGAP3 | 0.326 | 1.6E-02 |
| Q7Z3F1 | Integral membrane protein GPR155 GN=GPR155 | 0.320 | 3.1E-03 |
| Q9H2B2 | Synaptotagmin-4 GN=SYT4 | 0.320 | 3.3E-02 |
| Q7L1I2 | Synaptic vesicle glycoprotein 2B GN=SV2B | 0.318 | 1.2E-02 |
| O60268 | Uncharacterized protein KIAA0513 GN=KIAA0513 | 0.317 | 7.9E-03 |
| Q6U841 | Sodium-driven chloride bicarbonate exchanger GN=SLC4A10 | 0.317 | 3.8E-07 |
| Q9H0Q3 | FXYD domain-containing ion transport regulator 6 GN=FXYD6 | 0.314 | 2.0E-02 |
| P43146 | Netrin receptor DCC GN=DCC | 0.310 | 4.3E-04 |
| O94772 | Lymphocyte antigen 6H GN=LY6H | 0.310 | 1.4E-04 |
| O60939 | Sodium channel subunit beta-2 GN=SCN2B | 0.308 | 1.5E-03 |
| Q92797 | Symplekin GN=SYMPK | 0.301 | 3.0E-02 |
| Q5T848 | Probable G-protein coupled receptor 158 GN=GPR158 | 0.299 | 1.8E-04 |
| P08138 | Tumor necrosis factor receptor superfamily member 16 GN=NGFR | 0.298 | 3.2E-03 |
| Q7Z6B7 | SLIT-ROBO Rho GTPase-activating protein 1 GN=SRGAP1 | 0.295 | 1.4E-02 |
| Q9NWQ8 | Phosphoprotein associated with glycosphingolipid-enriched microdomains 1 GN=PAG1 | 0.294 | 8.1E-03 |
| O15118 | Niemann-Pick C1 protein GN=NPC1 | 0.293 | 2.7E-02 |
| Q8N7J2 | APC membrane recruitment protein 2 GN=AMER2 | 0.291 | 1.9E-03 |
| Q6UXK2 | Immunoglobulin superfamily containing leucine-rich repeat protein 2 GN=ISLR2 | 0.289 | 3.1E-04 |
| Q96GW7 | Brevican core protein GN=BCAN | 0.287 | 7.9E-03 |
| P26006 | Integrin alpha-3 GN=ITGA3 | 0.287 | 8.2E-04 |
| O43424 | Glutamate receptor ionotropic, delta-2 GN=GRID2 | 0.287 | 5.3E-04 |
| A2A2Y4 | FERM domain-containing protein 3 GN=FRMD3 | 0.283 | 4.7E-02 |
| Q9NZV1 | Cysteine-rich motor neuron 1 protein GN=CRIM1 | 0.282 | 5.9E-04 |
| P16389 | Potassium voltage-gated channel subfamily A member 2 GN=KCNA2 | 0.281 | 1.6E-02 |
| O60486 | Plexin-C1 GN=PLXNC1 | 0.276 | 2.1E-02 |
| Q99784 | Noelin GN=OLFM1 | 0.276 | 7.6E-03 |
| Q6IAA8 | Ragulator complex protein LAMTOR1 GN=LAMTOR1 | 0.271 | 3.8E-03 |
| P34903 | Gamma-aminobutyric acid receptor subunit alpha-3 GN=GABRA3 | 0.270 | 1.7E-02 |
| Q9UNL2 | Translocon-associated protein subunit gamma GN=SSR3 | -0.273 | 4.3E-02 |
| P46782 | 40S ribosomal protein S5 GN=RPS5 | -0.276 | 1.5E-02 |
| Q5VU97 | VWFA and cache domain-containing protein 1 GN=CACHD1 | -0.277 | 1.3E-03 |
| Q9Y5G3 | Protocadherin gamma-B1 GN=PCDHGB1 | -0.285 | 2.2E-04 |
| Q9Y5H3 | Protocadherin gamma-A10 GN=PCDHGA10 | -0.286 | 2.0E-05 |
| Q53HI1 | Protein unc-50 homolog GN=UNC50 | -0.287 | 3.8E-02 |
| Q96FZ5 | CKLF-like MARVEL tm domain-containing protein 7 GN=CMTM7 | -0.288 | 1.3E-02 |
| Q9ULU8 | Calcium-dependent secretion activator 1 GN=CADPS | -0.290 | 2.1E-02 |
| Q96RF0 | Sorting nexin-18 GN=SNX18 | -0.291 | 2.2E-02 |
| Q8IZU9 | Kin of IRRE-like protein 3 GN=KIRREL3 | -0.291 | 1.6E-04 |
| Q8WY07 | Cationic amino acid transporter 3 GN=SLC7A3 | -0.298 | 1.8E-02 |
| Q9UN71 | Protocadherin gamma-B4 GN=PCDHGB4 | -0.299 | 3.0E-04 |
| P19623 | Spermidine synthase GN=SRM | -0.299 | 4.0E-03 |
| Q8IXJ6 | NAD-dependent protein deacetylase sirtuin-2 GN=SIRT2 | -0.306 | 6.3E-03 |
| P55285 | Cadherin-6 GN=CDH6 | -0.307 | 5.5E-04 |
| Q9Y4C0 | Neurexin-3 GN=NRXN3 | -0.307 | 1.6E-03 |
| P54851 | Epithelial membrane protein 2 GN=EMP2 | -0.308 | 1.7E-02 |
| Q8NG11 | Tetraspanin-14 GN=TSPAN14 | -0.311 | 9.4E-03 |
| O43657 | Tetraspanin-6 GN=TSPAN6 | -0.311 | 8.1E-04 |
| P06756 | Integrin alpha-V GN=ITGAV | -0.317 | 1.4E-04 |
| Q14DG7 | Transmembrane protein 132B GN=TMEM132B | -0.325 | 1.5E-02 |
| P78504 | Protein jagged-1 GN=JAG1 | -0.326 | 6.3E-04 |
| P47972 | Neuronal pentraxin-2 GN=NPTX2 | -0.327 | 1.2E-03 |
| O14763 | TNF receptor superfamily member 10B GN=TNFRSF10B | -0.328 | 2.5E-03 |
| P0DKB5 | Trophoblast glycoprotein-like GN=TPBGL PE=4 | -0.329 | 2.5E-02 |
| Q9UBG0 | C-type mannose receptor 2 GN=MRC2 | -0.330 | 8.7E-03 |
| Q9UI15 | Transgelin-3 GN=TAGLN3 | -0.331 | 1.5E-02 |
| P04921 | Glycophorin-C GN=GYPC | -0.331 | 1.4E-04 |
| Q92870 | Amyloid beta A4 precursor protein-binding FbM2 GN=APBB2 | -0.332 | 7.4E-03 |
| Q9NZW5 | MAGUK p55 subfamily member 6 GN=MPP6 | -0.332 | 2.0E-04 |
| P51153 | Ras-related protein Rab-13 GN=RAB13 | -0.337 | 3.6E-02 |
| P42574 | Caspase-3 GN=CASP3 | -0.337 | 2.9E-04 |
| P60033 | CD81 antigen GN=CD81 | -0.348 | 8.8E-04 |
| Q75V66 | Anoctamin-5 GN=ANO5 | -0.352 | 3.8E-03 |
| P39019 | 40S ribosomal protein S19 GN=RPS19 | -0.356 | 4.8E-02 |
| Q15392 | Delta(24)-sterol reductase GN=DHCR24 | -0.357 | 2.6E-02 |
| Q13797 | Integrin alpha-9 GN=ITGA9 | -0.359 | 1.4E-07 |
| Q9ULK6 | RING finger protein 150 GN=RNF150 | -0.359 | 5.3E-03 |
| Q2VWP7 | Protogenin GN=PRTG | -0.363 | 9.1E-03 |
| Q9Y4D7 | Plexin-D1 GN=PLXND1 | -0.365 | 2.1E-04 |
| Q9H813 | Transmembrane protein 206 GN=TMEM206 | -0.367 | 1.0E-06 |
| Q9UN70 | Protocadherin gamma-C3 GN=PCDHGC3 | -0.368 | 3.1E-03 |
| Q14831 | Metabotropic glutamate receptor 7 GN=GRM7 | -0.368 | 6.7E-04 |
| O95628 | CCR4-NOT transcription complex subunit 4 GN=CNOT4 | -0.370 | 1.3E-02 |
| Q14517 | Protocadherin Fat 1 GN=FAT1 | -0.371 | 4.2E-05 |
| O00401 | Neural Wiskott-Aldrich syndrome protein GN=WASL | -0.374 | 3.9E-02 |
| P13611 | Versican core protein GN=VCAN | -0.379 | 3.4E-02 |
| P29317 | Ephrin type-A receptor 2 GN=EPHA2 | -0.380 | 1.7E-03 |
| O14522 | Receptor-type tyrosine-protein phosphatase T GN=PTPRT | -0.380 | 1.2E-03 |
| Q08431 | Lactadherin GN=MFGE8 | -0.381 | 5.1E-03 |
| P08582 | Melanotransferrin GN=MFI2 | -0.381 | 9.0E-04 |
| Q01973 | Tyrosine-protein kinase transmembrane receptor ROR1 GN=ROR1 | -0.383 | 2.9E-06 |
| A6NHL2 | Tubulin alpha chain-like 3 GN=TUBAL3 | -0.390 | 2.1E-02 |
| P24821 | Tenascin GN=TNC | -0.397 | 2.6E-02 |
| Q13464 | Rho-associated protein kinase 1 GN=ROCK1 | -0.413 | 2.2E-02 |
| Q13237 | cGMP-dependent protein kinase 2 GN=PRKG2 | -0.414 | 4.7E-02 |
| Q9Y625 | Glypican-6 GN=GPC6 | -0.419 | 1.4E-05 |
| P00533 | Epidermal growth factor receptor GN=EGFR | -0.444 | 7.0E-05 |
| Q12841 | Follistatin-related protein 1 GN=FSTL1 | -0.447 | 3.1E-03 |
| Q16698 | 2,4-dienoyl-CoA reductase, mitochondrial GN=DECR1 | -0.454 | 7.0E-04 |
| P29320 | Ephrin type-A receptor 3 GN=EPHA3 | -0.463 | 7.6E-04 |
| A6NEH6 | Transmembrane protein 247 GN=TMEM247 PE=4 | -0.471 | 1.8E-03 |
| Q494V2 | Coiled-coil domain-containing protein 37 GN=CCDC37 | -0.479 | 3.4E-02 |
| P58335 | Anthrax toxin receptor 2 GN=ANTXR2 | -0.481 | 2.1E-05 |
| P08648 | Integrin alpha-5 GN=ITGA5 | -0.499 | 5.4E-06 |
| P23471 | Receptor-type tyrosine-protein phosphatase zeta GN=PTPRZ1 | -0.538 | 1.5E-07 |
| P35221 | Catenin alpha-1 GN=CTNNA1 | -0.541 | 1.7E-06 |
| P61073 | C-X-C chemokine receptor type 4 GN=CXCR4 | -0.552 | 5.1E-04 |
| Q6UXZ4 | Netrin receptor UNC5D GN=UNC5D | -0.583 | 8.9E-04 |
| P56373 | P2X purinoceptor 3 GN=P2RX3 | -0.584 | 1.1E-05 |
| Q9UKU6 | Thyrotropin-releasing hormone-degrading ectoenzyme GN=TRHDE | -0.587 | 5.7E-05 |
| Q92563 | Testican-2 GN=SPOCK2 | -0.592 | 1.4E-02 |
| Q99653 | Calcineurin B homologous protein 1 GN=CHP1 | -0.595 | 1.6E-02 |
| Q9NZU0 | Leucine-rich repeat transmembrane protein FLRT3 GN=FLRT3 | -0.598 | 9.2E-03 |
| Q02297 | Pro-neuregulin-1, membrane-bound isoform GN=NRG1 | -0.619 | 1.2E-05 |
| O14495 | Lipid phosphate phosphohydrolase 3 GN=PPAP2B | -0.732 | 7.9E-06 |
| Q99996 | A-kinase anchor protein 9 GN=AKAP9 | -0.793 | 3.4E-05 |
| Q96A83 | Collagen alpha-1(XXVI) chain GN=COL26A1 | -0.620 | 1.3E-02 |
| O75487 | Glypican-4 GN=GPC4 | -0.915 | 2.5E-04 |

**Supplementary Table S4.** TaqMan Gene Expression Assay Probe Sets

| Gene | TaqMan Probe Set ID |
| --- | --- |
| GAPDH | Hs03929097_g1 |
| HEXA | Hs00166843_m1 |
| HEXB | Hs00166864_m1 |
| NANOG | Hs02387400_g1 |
| OCT4 | Hs04260367_gH |
| SYP | Hs00300531_m1 |
| MAP2 | Hs00258900_m1 |
| β3-tubulin | Hs00801390_s1 |
| LAMP-1 | Hs00174766_m1 |
| PSAP | HS01551096_m1 |
| CD63 | Hs01041238_g1 |
| BSN | Hs01109512_m1 |
| GPC4 | Hs00155059_m1 |
| NRG1 | Hs01101538_m1 |
| CNTN5 | Hs00544269_m1 |
| CNTNAP4 | Hs00369159_m1 |
| Syt1 | Hs00194572_m1 |

**Supplementary Table S5.** Purified lipids used as standards for PM glycan profiling

| Gene | Product code | Supplier |
| --- | --- | --- |
| Phosphatidylcholine | 840051C-25mg | Avanti Polar Lipids |
| Cholesterol | C8667-500MG | Sigma-Aldrich |
| Rhodamine-Phosphatidyl ethanolamine | 810150C-1MG | Avanti Polar Lipids |
| GM3 | 860058P-5MG | Avanti Polar Lipids |
| GM2 | G8397-1MG | Sigma-Aldrich |
| GM1a | 860065P-1MG | Avanti Polar Lipids |
| GD1a | 860055P-1MG | Avanti Polar Lipids |
| GD1b | 860056P-1MG | Avanti Polar Lipids |
| GT1b | 860059P-1MG | Avanti Polar Lipids |
| GQ1b | 860086P-1MG | Avanti Polar Lipids |

**Supplementary Methods**

***Proteomics - cell culture***

Cell lines for proteomic analysis were plated in triplicate to yield three independent biological replicates. Partially differentiated 3dpi i^3^Ns lines of SCRM, ΔHEXA-1, ΔHEXA-2, ΔHEXB-1 and ΔHEXB-2 cells were seeded at 2x10^6/ well in PLO coated plates for Whole-Cell Proteomics (WCP) and at 10x10^6 cells/ plate in PLO coated 10cm plates for Plasma Membrane Proteomics (PMP). Cells were left to differentiate until their 14dpi time-point with bi-weekly half media changes.

***Sample preparation for WCP***

Cells were washed three times with PBS and scaped into low-bind Eppendorf tubes. Cells were pelleted at 500g for 10 minutes, the PBS removed and snap frozen on LN2 and stored at -80°C until all samples were ready for simultaneous processing. Cell pellets were resuspended in 50 µL resuspension buffer (76 mM HEPES pH 7.55, 6 mM MgCl_2_, Benzonase (1400 U/mL) and 15 mM TCEP) by pipetting. Then, 18.75 µL of 20% lithium dodecyl sulfate was immediately added to the cell suspension using a low retention pipette tip (RPT, StarLab) and pipetted to mix. Nucleic acids were fragmented by 30 s on/30 s off sonication in a Bioruptor Pico sonicator (Diagenode) for 10 minutes at 4°C. Samples were then incubated for 15 minutes at 37 °C to ensure complete reduction. Samples were alkylated by adding 6 µL of 187.5 mM methyl methanethiosulfonate (final concentration 15 mM) and incubating at room temperature for 15 minutes. 5 µL aliquots of each sample were diluted 2x in water and compared to a standard curve of BSA in the same buffer using a reducing agent-compatible BCA assay (Thermo Fisher). 25 µg of each sample was taken and the volumes of each lysate were equalised using resuspension buffer with 5% lithium dodecyl sulfate.

To each sample a 10% volume of 12% phosphoric acid was added to acidify samples to ~pH 2, completing denaturation. 6x volumes of wash buffer (100 mM HEPES pH 7.1, 90% methanol) was then added and the resulting solution was loaded onto a S-trap (Protifi) using a positive pressure manifold (Tecan M10), adding not more than 150 µL of sample at a time (~80 PSI). In-house fabricated adaptors were used to permit the use of S-traps with the manifold. Samples were then washed 4x with 150 µL wash buffer. To remove any remaining wash buffer S-traps were centrifuged at 4000g for 2 minutes. To each S-trap, 30 uL of digestion solution (50 mM HEPES pH 8, 0.1% sodium deoxycholate) containing 1 µg Trypsin/lysC mix (Promega) was added. S-Traps were then loosely capped and placed in low adhesion 1.5 mL microfuge tubes in a ThermoMixer C (Eppendorf) with a heated lid and incubated at 37 °C for 6 hours. Where digestion was carried out overnight the Thermomixer was set to 4 °C after 6 hours. Peptides were recovered by adding 40 µL digestion buffer to each trap and incubating at room temperature for 15 minutes before slowly eluting with positive pressure (2-3 PSI). Traps were subsequently eluted with 40 µL 0.2% formic acid and 40 µL 0.2% formic acid, 50% acetonitrile in the same manner. Eluted samples were then dried in a vacuum centrifuge equipped with a cold trap prior to TMT labelling.

***Sample preparation for PMP-MS***

Sialic acid moieties present on the extracellular side of PM proteins were oxidised to produce an aldehyde group using Sodium Periodate, followed by an analine-catalysed oxime ligation to aminooxy-biotin. Cells prepared for PMP were washed three times with 5ml of ice-cold PBS pH 7.4 and then incubated under 5ml of biotinylation mix (1mM Na Periodate, 100µM aminooxy-biotin, 10mM aniline in ice-cold PBS pH 6.7) and wrapped in foil with rocking, at 4°C for 30 minutes. After biotin labelling, the reaction was quenched with 5 ml of 2 mM glycerol, the biotinylation/quenching reaction was then removed, and the cells washed 3x with PBS pH 7.4. All PBS was removed, and cells were scraped into a low bind Eppendorf in 500 μl of lysis buffer (10mM Tris pH 7.4, 1% Triton X-100, 150 mM NaCl, 5 mM EDTA and 1 protein inhibitor tab per 50 ml).

Samples in lysis buffer were lysed through end over end rotation for 1.5 hours at 4°C. Lysates were clarified with centrifugation at 20,000g for 10 minutes. The supernatant was then removed to a fresh low-bind Eppendorf tube and snap frozen on LN2 and stored at -80°C until all samples were ready for simultaneous enrichment steps. Frozen samples were thawed, and a BCA assay was performed. Samples were normalised with lysis buffer to the lowest concentration to ensure the same mass of protein went into subsequent enrichment steps. 50ul of Neutravidin bead slurry per sample was washed with 1ml of lysis buffer four times (including resuspension, centrifugation 500g for 5 minutes, removal of liquid, repeat). Normalised samples were then added to neutravidin beads and incubated with end over end rotation for 2.75 hours at 4°C.

Beads with enriched samples were washed using a vacuum manifold and SnapCap filter columns (Pierce). Bead/sample slurry was moved into the snapcap columns and allowed to drain. Each sample was then washed 20x with 400 μl of lysis buffer, followed by 20x washes with 0.5% SDS in PBS pH 7.4, followed by 10x washes with Urea buffer (6M urea, 1M TEAB (Thermo), pH 8.5). Columns were removed, capped to prevent liquid loss and the beads were incubated with 400 μl of reduction/alkylation solution (10 mM TCEP (Thermo), 20 mM Iodoacetamide, in Urea buffer), shaking at 850 rpm for 30 minutes in the dark. Samples were returned to the vacuum manifold and the beads drained before a further 10x washed in urea buffer. Columns were again capped, and the beads resuspended in 400 μl of 50 mM TEAB and removed to a low adhesion Eppendorf tube. The columns were rinsed again 2x with 400 μl of 50 mM TEAB and the washes combined with the sample. Beads were pelleted gently at 500 g for 2 minutes and all supernatant was removed. Beads were then resuspended in 50 μl of 50 mM TEAB + 0.5 μg of trypsin (MS grade, Pierce) (trypsin stocks made up at 1 μg/μl in 50 mM acetic acid) and incubated at 850 rpm and 37°C, overnight. The following day, the beads were pelleted at 500g for 5 minutes and the supernatant removed and stored. The beads were washed with a further 40 μl of 50 mM TEAB and pelleted again, the wash supernatant was combined with the sample and then samples were dried in a vacuum centrifuge before storage at -20°C awaiting TMT labelling.

***TMT labelling and clean-up***

Dried samples were resuspended in 21 µL 100 mM TEAB pH 8.5. After warming to room temperature, 0.5 µg TMTpro/0.2 µg TMT reagents (Thermo Fisher) were resuspended in 9 µL anhydrous acetonitrile which was added to the respective samples and incubated at room temperature for 1 h. A 3 µL aliquot of each sample was taken and pooled to check TMT labelling efficiency and equality of loading by liquid chromatography-mass spectrometry (LC-MS). Samples were stored at -80 °C in the interim. After checking each sample was at least 98% TMT labelled, total reporter ion intensities were used to normalise the pooling of the remaining samples such that the final pool should be as close to a 1:1 ratio of total peptide content between samples as possible. This final pool was then dried in a vacuum centrifuge. Whole cell proteomics samples were acidified to a final 0.1% trifluoracetic acid (~200 µL volume) and formic acid was added until the sodium deoxycholate visibly precipitated. 4 volumes of ethyl acetate were then added and the sample vortexed vigorously for 10 s. The sample was then centrifuged at 15,000g for 5 min at room temperature to effect phase separation. A gel loading pipette tip was used to withdraw the lower (aqueous) phase to a fresh low adhesion microfuge tube. If any obvious sodium deoxycholate contamination remained, the two-phase extraction with ethyl acetate was repeated. The sample was then partially dried in a vacuum centrifuge. Whole cell and PMP samples were brought up to a final volume of 1 mL with 0.1% trifluoracetic acid. Formic acid (FA) was added until the pH was < 2, confirmed by spotting onto pH paper. The sample was then cleaned up by solid-phase extraction using a 50 mg tC18 SepPak cartridge (Waters) and a positive pressure manifold. The cartridge was wetted with 1 mL 100% Methanol followed by 1 mL acetonitrile, equilibrated with 1 mL 0.1% trifluoracetic acid and the sample was loaded slowly. The sample was passed twice over the cartridge. The cartridge was washed 3x with 1 mL 0.1% trifluoracetic acid before eluting sequentially with 250 µL 40% acetonitrile, 70% acetonitrile and 80% acetonitrile and dried in a vacuum centrifuge.

***Mass spectrometry***

Mass spectrometry data was acquired using an Orbitrap Lumos as previously described [1]. An Ultimate 3000 RSLC nano UHPLC equipped with a 300 µm ID x 5 mm Acclaim PepMap µ-Precolumn (Thermo Fisher Scientific) and a 75 µm ID x 50 cm 2.1 µm particle Acclaim PepMap RSLC analytical column was used. Loading solvent was 0.1% FA, analytical solvent A: 0.1% FA and B: 80% MeCN + 0.1% FA. All separations were carried out at 40°C. Samples were loaded at 5 µL/minute for 5 minutes in loading solvent before beginning the analytical gradient. The following gradients were used: 3-7% B over 4 minutes, 7-37% B over 173 minutes, followed by a 4-minute wash at 95% B and equilibration at 3% B for 15 minutes or 3-7% B over 3 minutes, 7-37% B over 116 minutes, followed by a 4-minute wash at 95% B and equilibration at 3% B for 15 minutes. Each analysis used a MultiNotch MS3-based TMT method [2]. The following settings were used: MS1: 380-1500 Th, 120,000 Resolution, 2x105 automatic gain control (AGC) target, 50 ms maximum injection time. MS2: Quadrupole isolation at an isolation width of m/z 0.7, CID fragmentation (normalised collision energy (NCE) 34) with ion trap scanning in turbo mode with 1.5x104 AGC target and 120 ms maximum injection time. MS3: In Synchronous Precursor Selection mode the top 10 MS2 ions were selected for HCD fragmentation (NCE 45) and scanned in the Orbitrap at 60,000 resolution with an AGC target of 1x105 and a maximum accumulation time of 150 ms. Ions were not accumulated for all parallelisable time. The entire MS/MS/MS cycle had a target time of 3 s.

***Analysis of mass spectrometry data***

Data was processed in Proteome Discoverer 2.2 using the Mascot search engine and a Uniprot human database (downloaded 11/01/2021). The results were imported into Perseus [3] as a tab delimited text file. In Perseus the grouped reporter abundances were log_2_ transformed, data filtered to a minimum of 3 valid values and missing values replaced from a normal distribution. The replicates were grouped, and volcano plots generated, with significantly changed proteins determined using a two-sample t-test with an FDR cutoff of <0.05. Differences in protein abundance and adjusted P-value were used to ascribe significance and to generate volcano plots in GraphPad Prism.

The Database for Annotation, Visualisation and Integrated Discovery (DAVID) was used for functional annotation and enrichment analysis of proteomic datasets [4]. Proteins that were significantly changed (p<0.05) in abundance by >20% compared to SCRM controls and with ≥2 PSMs were searched against a background reference list of all proteins quantified in the respective dataset with ≥2 PSMs. For functional enrichment analysis of WCP, only proteins increased in abundance were selected for analysis against a background list as defined above. Cellular Component GO_Direct terms were then selected based on FDR.
